# Tissue-specific profiling reveals distinctive regulatory architectures for ubiquitous, germline and somatic genes

**DOI:** 10.1101/2020.02.20.958579

**Authors:** Jacques Serizay, Yan Dong, Jürgen Jänes, Michael Chesney, Chiara Cerrato, Julie Ahringer

## Abstract

Despite increasingly detailed knowledge of gene expression patterns, the regulatory architectures that drive them are not well understood. To address this, we compared transcriptional and regulatory element activities across five adult tissues of *C. elegans*, covering ∼90% of cells, and defined regulatory grammars associated with ubiquitous, germline and somatic tissue-specific gene expression patterns. We find architectural features that distinguish two major promoter types. Germline-specific and ubiquitously-active promoters have well positioned +1 and −1 nucleosomes associated with a periodic 10-bp WW signal. Somatic tissue-specific promoters lack these features, have wider nucleosome depleted regions, and are more enriched for core promoter elements, which surprisingly differ between tissues. A 10-bp periodic WW signal is also associated with +1 nucleosomes of ubiquitous promoters in fly and zebrafish but is not detected in mouse and human. Our results demonstrate fundamental differences in regulatory architectures of germline-active and somatic tissue-specific genes and provide a key resource for future studies.

## Introduction

Cell-type specific transcription regulation underlies production of the myriad of different cells generated during development. Regulatory elements (*i.e.* promoters and enhancers) are key sequences that direct appropriate spatio-temporal gene expression patterns, and they can have diverse activities, ranging from ubiquitous to highly cell-type specific (Cusanovich et al. 2018; Liu et al. 2019; Smith et al. 2007; Andersson and Sandelin 2019). The composition, activity and arrangement of regulatory elements define the regulatory grammar that controls patterns of gene transcription across development (Abhijeet Rajendra Sonawane et al. 2017; Heinz et al. 2015; Levine 2010; Ong and Corces 2011; Spitz and M Furlong 2012) and mutation or perturbation of their spatial organization can lead to pathologies (Lupiáñez et al. 2016).

Previous studies have provided important and increasingly detailed knowledge of features of transcription regulation in eukaryotes. Different regulatory architectures have been observed, ranging from single promoters to complex structures involving multiple regulatory elements, which can operate redundantly, hierarchically, additively or synergistically (Osterwalder et al. 2018; Herr 1993; Bahr et al. 2018; Guerrero et al. 2010; Davuluri et al. 2008; Whyte et al. 2013). Work on human cells suggests that housekeeping genes are primarily regulated by a single core promoter whereas tissue-specific genes rely on additional regulatory elements (Ernst et al. 2011). Moreover, differences in sequence features, patterns of transcription initiation and nucleosome arrangement characterize promoters with different activities (Lenhard et al. 2012; Haberle and Lenhard 2016). Yet, cell type specific differences are still not well understood. More comprehensive genome-wide *in vivo* studies of regulatory grammar would directly address how specific gene expression patterns in different tissues are achieved and whether expression is governed by distinct regulatory architectures. Towards this end, we profiled and compared nuclear transcriptomes and chromatin accessibility in sorted *C. elegans* adult tissues. Through analyses of these rich datasets, we uncover shared distinct features of ubiquitous and tissue-specific regulatory architectures.

## Results

### Tissue-specific profiling of chromatin accessibility and gene expression in adult *C. elegans* tissues

To investigate the regulatory chromatin of different cell types and how it relates to gene expression, we developed a procedure to isolate nuclei from individual tissues in *C. elegans*. We expressed GFP tags on the outside of the nuclear envelope using tissue-specific promoters and isolated labelled nuclei using fluorescent activated nuclear sorting (Fig. 1A, Supplemental Fig. S1, S2; see Methods). Applying this methodology to young adult animals using promoters active in the five major tissues of *C. elegans,* we could obtain nuclei of high purity (97.4% ± 1.27 SD) from the germ line and four somatic tissues (muscle, hypodermis, intestine, and neurons; Supplemental Table S1). This covers ∼90% of cells in young adults but does not include the pharynx, glia, or somatic gonad.

**Figure 1.**
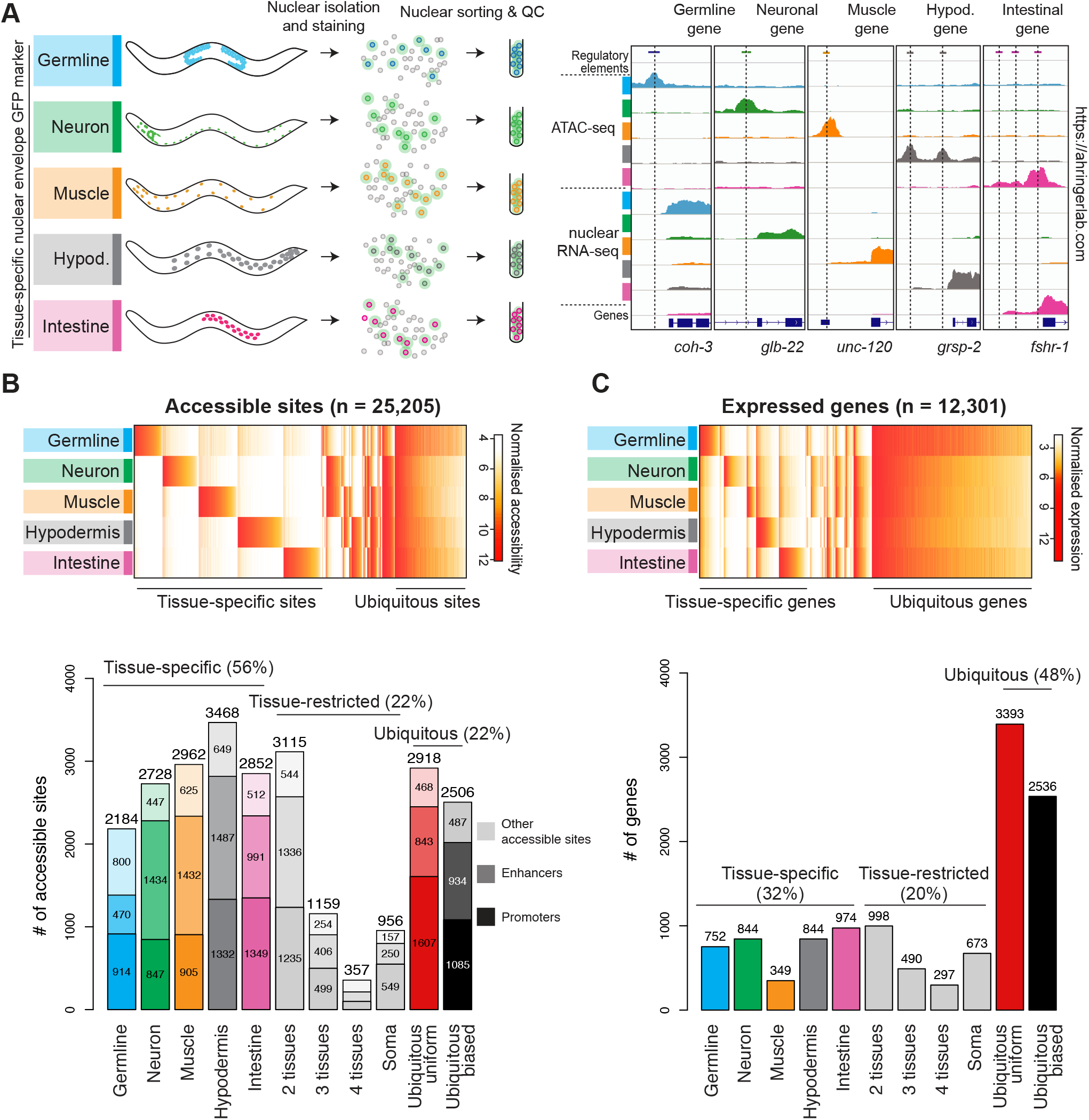
Tissue-specific profiling of chromatin accessibility and gene expression in adult *C. elegans* tissues. (A) Procedure to perform tissue-specific nuclear RNA-seq and ATAC-seq experiments in young adults. Representative results at known tissue-specific loci are shown on the right. (B) Top: heatmap of normalised accessibility (log2 RPM) for 25,205 classified sites. Bottom: classification of the accessible sites into tissue-specific, tissue-restricted or ubiquitous classes. Protein-coding promoters are in dark colors, putative enhancers are lighter and other accessible sites (*e.g.* non-coding promoters, unassigned promoters, other elements) are lightest. (C) Top: heatmap of normalised gene expression (log2 TPM) for 12,301 classified protein-coding genes. Bottom: classification of genes into tissue-specific, tissue-restricted or ubiquitous classes. See methods for classification procedure. Unclassified sites and genes are not shown.

We previously defined 42,245 accessible elements across *C. elegans* development and ageing in whole animals using ATAC-seq, and annotated promoters, putative enhancers, and other accessible sites based nuclear RNA-seq patterns (Jänes et al. 2018). To classify the adult tissue-specificity of gene expression and accessibility, and to identify new elements, we carried out ATAC-seq and nuclear RNA-seq on sorted nuclei from the five tissues. Biological replicates were highly concordant (Supplemental Fig. S2D-E) and known tissue-specific loci showed expected activities (Fig. 1A). We identified 5,269 additional accessible elements through these new data, bringing the total to 47,514, with 11,806 genes now having at least one high-confidence promoter. We then classified the tissue-specificity of element accessibility and gene expression using a set of conservative rules (see Methods). Excluding elements and genes with low signal in all assessed samples, 25,205 (53%) of the 47,514 accessible sites, and 12,301 (61%) of the 20,222 protein-coding genes were classified (Fig. 1B-C and Supplemental Table S2).

We observed that the chromatin accessibility of regulatory elements is largely tissue-specific. The majority of regulatory elements (56%) are accessible in only a single tissue, with the rest having tissue restricted (22%) or ubiquitous accessibility (22%); the latter were split into those with relatively uniform accessibility (<3-fold difference between any two tissues) and those with biased accessibility (Fig. 1B). For gene expression, the largest class of genes (48%) had ubiquitous expression, with the remainder having tissue-specific (32%) or tissue-restricted (20%) expression (Fig. 1C). The gene expression classification showed good overlap with previously published annotations ((Cao et al. 2017; Kaletsky et al. 2018); Supplemental Fig. S2G and S2H). We observed that the nuclear RNA datasets have minor contamination by bulk cytoplasmic RNA released during nuclear isolation, which resulted in tissue-specific genes with high expression (e.g, muscle myosin gene *unc-54*) being classified as ubiquitous-biased. Hereafter, when studying ubiquitous genes and elements, we specifically focus on the ubiquitous-uniform class and for simplicity refer to them as “ubiquitous”.

The data provide a comprehensive view of chromatin accessibility and transcriptional landscapes in the five major adult tissues. We have created the *C. elegans* regulatory atlas (RegAtlas, https://ahringerlab.com/) to facilitate access and analyses of these new tissue-specific and previous development datasets (Jänes et al. 2018). Below, we analyse features of genes and regulatory elements active in different tissues.

### Germline-active and soma-restricted genes have distinctive regulatory architectures

To investigate whether general rules could be discerned that govern different types of spatial expression patterns, we focused on genes with ubiquitous or tissue-specific expression and compared the number, type and arrangement of regulatory elements associated with genes from each class.

As expected, most (77%) ubiquitously expressed genes with at least one classified promoter are associated with a ubiquitously active promoter (Supplemental Table S2). We observed that half (54%) of the ubiquitous genes have just a single promoter, whereas 16% have a relatively complex regulatory architecture containing three or more promoters (Fig. 2A). To explore these differences, we separated ubiquitously expressed genes into groups based on promoter number (one, two, three or more). We observed that single promoter genes are enriched for functions in basic cellular processes whereas those with three or more promoters are enriched for developmental functions (Fig. 2B). Multi-promoter ubiquitous genes also have more enhancers than single promoter genes, are more often controlled by unidirectional promoters, and have more and longer introns (Fig. 2C-F).

**Figure 2.**
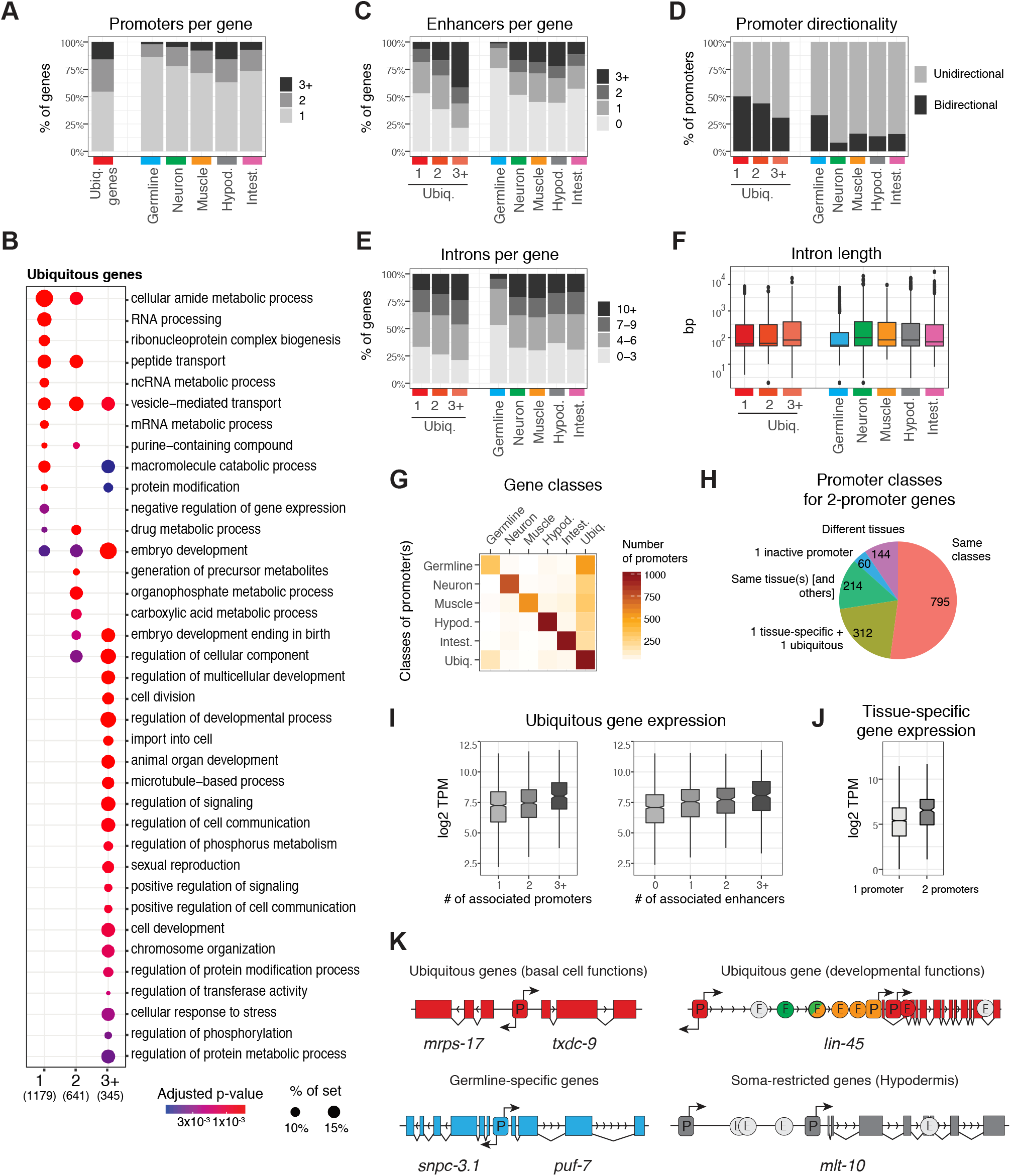
Regulatory architectures of ubiquitous, germline and soma-restricted genes have distinctive features. (A) Percentage of genes with one, two, or three or more promoters for each gene class. (B) GO terms from Biological Process ontology enriched in ubiquitous genes with one, two, or three or more annotated promoters. (C) Percentage of genes with zero, one, two, or three or more enhancers associated with genes of each expression class. Only genes with at least one annotated promoter are considered. (D) Percentage of unidirectional or bidirectional protein-coding promoters for each gene class. (E) Percentage of genes with the indicated number of introns for each gene class. (F) Intron length for each gene class. (G) Classes of promoters associated with genes of each expression class. Only the major promoter classes are displayed. See Supplemental Table S2 for all results. (H) Concordance of promoter classes for genes with two promoters. (I) Gene expression levels in whole young adults for ubiquitous genes with one, two, or three or more promoters (left), or zero, one, two, or three or more enhancers (right). (J) Gene expression levels of tissue-specific genes with one promoter or two promoters specifically active in the same tissue. (K) Left: examples of the simple regulatory architecture shared by ubiquitous genes and germline-specific genes. Right: examples of more complex architectures found at developmental ubiquitous genes (*e.g. lin-45*) or somatic tissue-specific genes (*e.g. mlt-10*).

As for ubiquitous genes, tissue-specific genes are generally associated with one or more promoters specific for the corresponding tissue (78%, Fig. 2G). Intriguingly, we observed that a group of genes with germline-specific expression have ubiquitously accessible promoters (Fig. 2G). We found that these genes are enriched for being targets of the repressive Rb/DREAM complex (13-fold enrichment, p-value = 5e-13) and in line with this, they are enriched for cell cycle and cell division functions (Supplemental Fig. S3A-B; (Latorre et al. 2015; Goetsch et al. 2017). This suggests that the predominantly germline expression of these genes at the young adult stage is achieved via their silencing in somatic tissues (Petrella et al. 2011; Wu et al. 2012).

Comparing the different tissue-specific classes, we found that germline-specific genes show extensive differences compared with somatic genes. First, germline-specific genes have fewer promoters and enhancers than somatic genes (Fig. 2A, 2C); 65% of germline-specific genes with at least one classified promoter have a single promoter and no associated enhancer, compared to 38% of somatic genes. The promoters of germline genes are more often bidirectional than those of somatic genes (Fig. 2D), and germline genes also have fewer and shorter introns, similar to ubiquitously expressed single promoter genes (Fig. 2E-F).

A significant fraction of expressed genes with at least one annotated promoter (33%) have more than one promoter, and alternative promoters are frequently active in the same tissue (Fig. 2H and Supplemental Fig. S3C; Supplemental Table S2), suggesting that alternative promoters may play a role in the regulation of expression levels. To investigate this, we examined the relationship between the number of regulatory elements and gene expression level. Among ubiquitously expressed genes, we found that the number of promoters and enhancers was positively correlated with gene expression (Fig. 2I). Similarly, tissue-specific genes with two tissue specific promoters have higher gene expression levels than those with only one (Fig. 2J). We also note that 15% of the ubiquitously expressed genes with two promoters have one tissue-specific promoter in addition to a ubiquitously active one, which could be a mechanism to increase gene expression specifically in a particular tissue (Supplemental Fig. S3D). These results suggest that an important but often overlooked role of regulatory elements is to augment gene expression rather than being necessary for its expression *per se*. This could explain some cases where deletion of an individual regulatory element does not have an obvious effect on gene expression despite its having activity in a transgenic assay (Dukler et al. 2016; Catarino and Stark 2018).

To summarize, we found that the regulatory architecture of genes is related to their function and expression pattern (Fig. 2L). Ubiquitous genes required for fundamental cellular processes and germline-specific genes tend to have a simple architecture consisting of a single promoter that is often bidirectional. In contrast, ubiquitous genes with functions associated with multicellular life often have a more complex architecture of multiple regulatory elements that can have diverse tissue specificity. Somatic tissue-specific genes usually have one or more regulatory elements accessible only in the matching tissue. Finally, the positive relationship between gene expression and the number of regulatory elements supports a role in modulating the level of gene expression.

### Ubiquitous and germline-specific promoters have a stereotypical architecture with well-positioned nucleosomes

The tissue-specific differences in gene regulatory architectures prompted us to investigate whether differences also occurred at the level of promoters. Comparing accessibility patterns of different classes of promoters, we observed that germline-specific and ubiquitously active promoters were flanked by regions of increased accessibility and associated with more nucleosome-sized ATAC-seq fragments, suggesting the presence of well positioned nucleosomes (Supplemental Fig. S4A-B). The flanking ATAC-seq signal at germline promoters was also present in proliferative stages (L1 and L3 larval larvae), indicating that it is not simply a characteristic of adult germline nuclei undergoing meiosis (Supplemental Fig. S4C).

To investigate this potential signature of positioned nucleosomes, we used ATAC-seq fragment density plots (also known as “V-plots”, (Henikoff et al. 2011) to visualize the distribution of fragment lengths relative to the distance to promoter center. Over promoters flanked by positioned −1 and +1 nucleosomes, V-plots show stereotypical patterns with a central concentration of small fragments at the nucleosome-depleted region (NDR) and larger fragments over +1/-1 nucleosomes on either side of the NDR (Henikoff et al. 2011); Supplemental Fig. S5A). In line with this, a signature of −1 and +1 nucleosomes is readily apparent at ubiquitous promoters in all tissues, as well as at germline-specific promoters (Fig. 3A). However, somatic tissue-specific promoters lack this signature of well-positioned +1/-1 nucleosomes (Fig. 3A, Supplemental Fig. S5B-D), regardless of the level of expression of the associated gene (Supplemental Fig. S5E).

**Figure 3.**
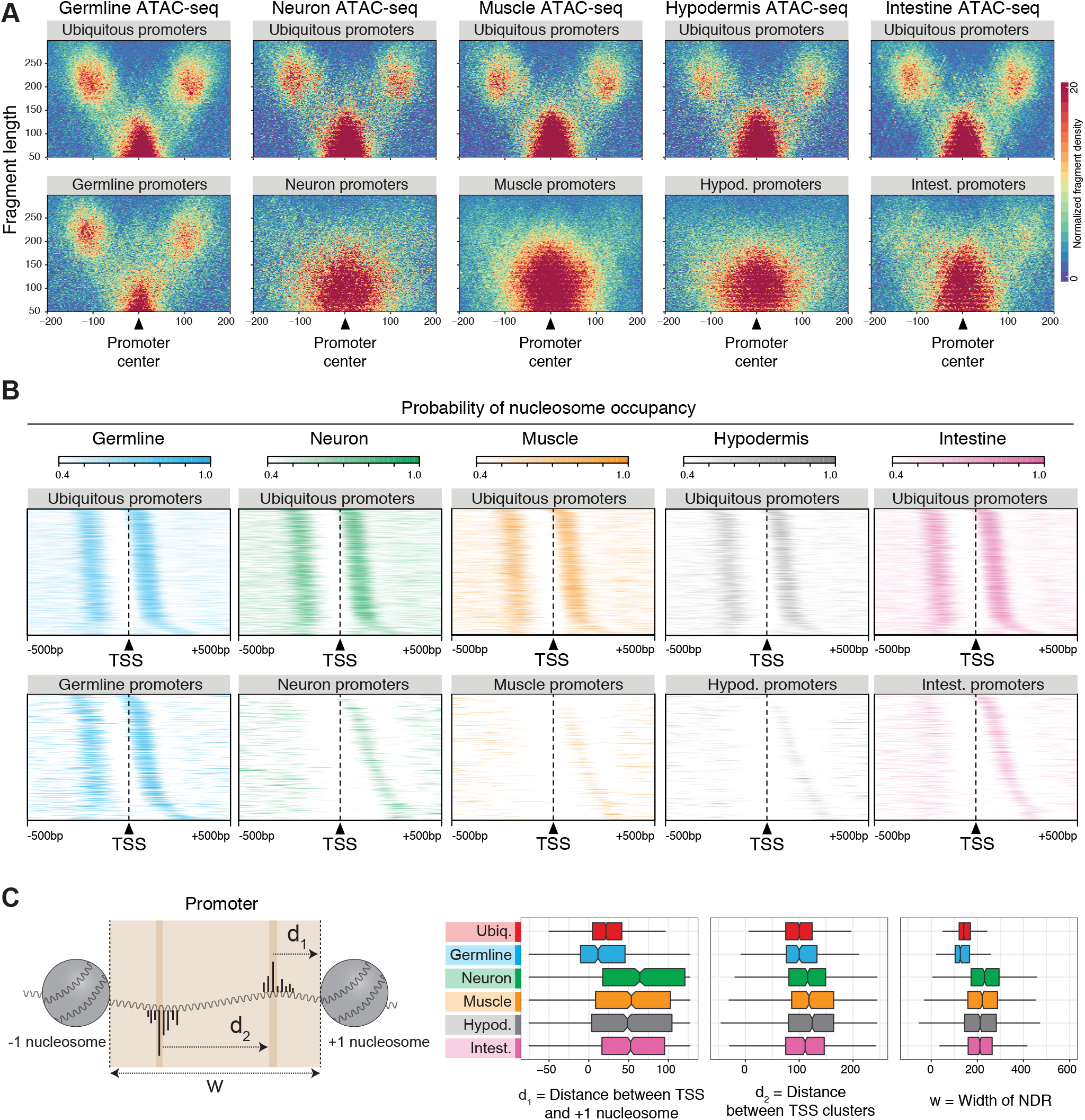
Ubiquitous and germline-specific promoters have a stereotypical architecture with well-positioned nucleosomes. (A) ATAC-seq fragment density plots (also known as “V-plots”) over different classes of promoters. The x axis represents the distance between the fragment midpoint and the promoter center. The y axis represents ATAC-seq fragment length. The color scale indicates the normalized density of ATAC-seq fragments. (B) Tissue-specific nucleosome occupancy probability over different classes of promoters aligned at their TSS. Only promoters with experimentally defined forward and reverse TSSs are considered. Rows are ordered by the distance between TSS and +1 nucleosome. (C) Left: schematic of the distance metrics measured in promoters: d_1_, distance between the mode TSS and the +1 nucleosome edge; d_2,_ distance between modes of divergent TSSs within the same promoter; w, width of the nucleosome-depleted region (NDR). Right: d_1_, d_2_ and w distance metrics for different classes of promoters. The metrics for ubiquitous promoters were measured using nucleosome occupancy probability track derived from whole young adult ATAC-seq data (Jänes et al. 2018).

To explore this further, we used the ATAC-seq data to compute nucleosome occupancy probability profiles as in (Schep et al. 2015). This revealed a high probability of +1 and −1 nucleosome occupancy at consistent positions relative to TSSs of ubiquitous and germline-specific promoters (Fig. 3B and Supplemental Fig. S4D). In contrast, somatic tissue-specific promoters were characterized by lower −1 and +1 nucleosome occupancy and a larger range of nucleosome positions relative to TSSs (Fig. 3B and Supplemental Fig. S4D). We find that the 5’ edges of +1 nucleosomes at ubiquitous and germline-specific promoters have narrow distributions relative to TSSs, with median distances of 22 bp for ubiquitous promoters and 12 bp for germline-specific promoters. In contrast, +1 nucleosomes at somatic tissue-specific promoters have much wider distributions and larger median distances (Fig. 3B-C). We also observed that NDR widths are smaller and divergent promoter TSSs are closer together for ubiquitous and germline-specific promoters compared with somatic tissue-specific promoters (Fig. 3C and Supplemental Fig. S4D). Of note, the median NDR widths at ubiquitous and germline-specific promoters are 140 bp and 125 bp, which would be too short to accommodate a nucleosome.

To identify sequence features that may be responsible for these differences, we carried out motif analyses. We observed that ubiquitous and germline-specific promoters share a T-rich motif with 10 bp spacing that was not present at somatic tissue-specific promoters (Fig. 4A). Previous studies mostly performed *in vitro* or in yeast have implicated 10-bp WW (W = A/T) periodicity in nucleosome positioning and observed SS periodicity in antiphase with WW (Satchwell et al. 1986; Ioshikhes et al. 1996; Wang and Widom 2005; Segal et al. 2006; Johnson et al. 2006; Mavrich et al. 2008a; Field et al. 2008; Struhl and Segal 2013).

**Figure 4.**
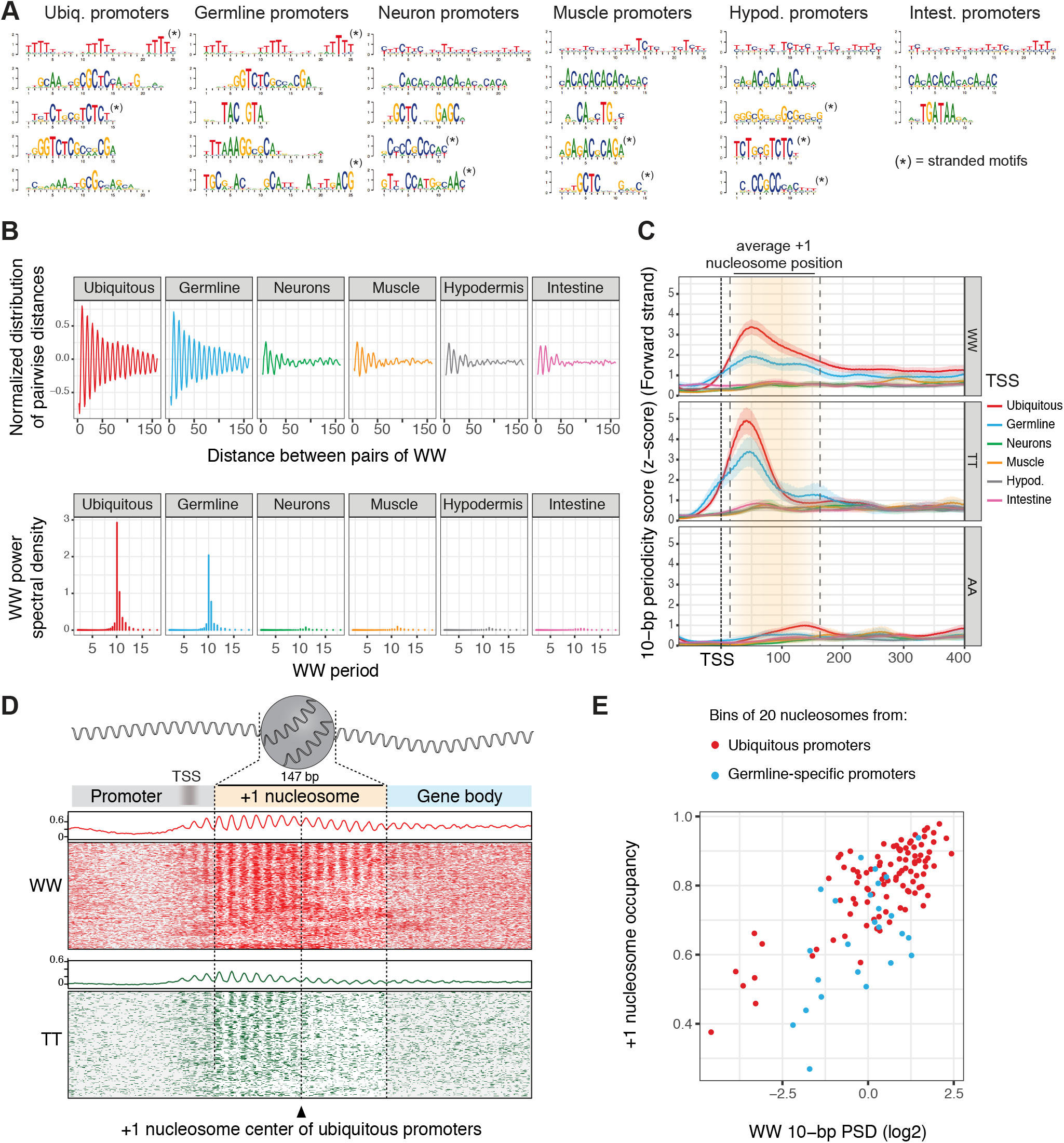
Ubiquitous and germline-specific promoters have strong 10-bp WW periodicity correlated with nucleosomes. (A) Motifs enriched in different classes of promoters. Sequences from −75 to +105 bp around the promoter centers were considered. (B) Top: Normalized distribution of pairwise distances between WW dinucleotides found in the sequences from −50 bp to +300 bp relative to TSSs, for different classes of promoters. Bottom: associated WW power spectral densities (PSDs). (C) Metaplots of WW, TT and AA 10-bp periodicity scores at different classes of promoters, aligned at TSSs. The +1 nucleosome position observed at ubiquitous and germline promoters (∼ 20-167 bp downstream of the TSS) is displayed by the shaded orange area delimited by dotted lines. (D) WW (red) and TT (green) dinucleotide occurrences observed at +1 nucleosomes of ubiquitous promoters (400 bp window centered at nucleosome dyads). Rows were shifted up to 5 bp to highlight the phased 10-bp periodic patterns. Summed dinucleotide occurrences are represented on top of each heatmap by a line plot. The average TSS positions of ubiquitous promoters (∼20 bp upstream of the +1 nucleosome edge) are displayed by the shaded gray area. (E) Correlation between +1 nucleosome occupancy and 10-bp WW periodicity in ubiquitous and germline-specific promoters. +1 nucleosomes were binned by their nucleosome occupancy score and the overall 10-bp WW periodicity was assessed in each bin (∼ twenty 200-bp long nucleosomal sequences centered at nucleosome dyads). The y axis represents the average nucleosome occupancy in each bin.

To investigate whether the T-rich motif we identified was part of a larger WW periodic signal involved in +1 nucleosome positioning at ubiquitous and germline-specific promoters in *C. elegans*, we measured WW dinucleotide periodicity from −50 bp to +300 bp relative to TSSs. We observed that ubiquitous and germline-specific promoter regions harbor a strong 10-bp periodic WW signal that extends for more than 150 bp and that the periodicity signal coincides with +1 nucleosome position (Fig. 4B-D, Supplemental Fig. S6A). Furthermore, at these promoters we found that 10-bp WW periodicity strength is correlated with +1 nucleosome occupancy (Fig. 4E). In contrast, the 10-bp periodic WW signal was not detected at somatic-tissue-specific promoters, in line with the absence of positioned +1 nucleosomes at these promoters (Fig. 4B-C, Supplemental Fig. S6A-B). Therefore, an extended 10-bp periodic WW signal specific to ubiquitously active and germline active promoters is associated with nucleosome position and occupancy.

Examining the contribution of different dinucleotides to the WW signal, we found that TT periodicity peaks in the 5’ region of the +1 nucleosome of ubiquitous and germline-specific promoters, ∼50 bp downstream of the TSS, and makes a larger contribution than other dinucleotides (Fig. 4C-D, Supplemental Fig. S6A). A weaker AA periodic signal peaks at the 3’ edge of the nucleosome (Fig. 4C-D, Supplemental Fig. S6A), and AT and TA dinucleotides do not show any robust periodic signal (Supplemental Fig. S6A). We note that a 10-bp SS periodicity antiphase with WW is also present at ubiquitous and germline-specific promoters (Supplemental Fig. S6A-B).

The strength of the 10-bp periodic WW signal is similar at +1 nucleosomes of bidirectional and unidirectional ubiquitous promoters (Supplemental Fig. S6D). Periodicity is also present at −1 nucleosomes of the unidirectional promoters, although the signal is weaker (Supplemental Fig. S6D). We also note that WW periodicity strength differs among ubiquitous promoters of ubiquitous genes. WW periodicity is stronger at single promoter genes, which are enriched for basal cell functions, compared to ubiquitous promoters of genes with three or more promoters, which are enriched for developmental functions (Fig. 2B and Supplemental Fig. S6D).

We next investigated the tissue-specificity and position of other promoter elements. The Inr initiator sequence, the Sp1 motif and the TATA-box are three well-known core promoter elements that have been previously observed in *C. elegans* promoters (Chen et al. 2013; Saito et al. 2013). Inr motifs were detected in all promoter classes, however, somatic tissue-specific promoters showed higher enrichment than ubiquitous and germ-line specific promoters (Supplemental Fig. S7A). We further observed that the Sp1 and TATA box motifs were both predominantly associated with somatic tissue-specific promoters, with striking and unexpected tissue biases (Supplemental Fig. S7A). The Sp1 motif, peaking at −45bp from the TSS, is enriched at neural, muscle and hypodermal promoters but not at intestinal promoters, whereas the TATA-box motif was predominantly found at hypodermal and intestinal promoters. We also observed that somatic tissue-specific promoters share repeated dinucleotide composition biases not found in ubiquitous or germline-specific promoters (Fig. 4A and Supplemental Fig. S7A).

The *de novo* motif analyses also uncovered motifs associated with promoters active in single tissues (Fig. 4A, Supplemental Fig. S7). For example, as expected, many intestinal promoters harbor a GATA motif, while the HLH-1 motif is found specifically at muscle promoters (Supplemental Fig. S7; (McGhee et al. 2007; Chen et al. 1994). These motifs and others have peak positions within the NDR, often ∼45 bp upstream of the TSS (Supplemental Fig. S7). Thus, there are tissue-specific differences in both core promoter elements and TF binding motifs.

In summary, our results uncover two largely different types of promoter architecture. Ubiquitous and germline-specific promoters have well-positioned +1/-1 nucleosomes that are highly associated with a periodic 10-bp WW signal and stereotypically positioned with the 5’ edge ∼20 bp downstream of the TSS, and they have relatively short nucleosome depleted regions. In contrast, +1 nucleosomes of somatic tissue-specific promoters have low occupancy and inconsistent positioning relative to TSSs, and nucleosome depleted regions are wider. In addition, core promoter and transcription factor motifs show strong tissue biases.

### 10-bp WW periodicity at ubiquitous promoters is a feature of non-mammalian genomes

We next asked whether a 10-bp periodic WW signal is a feature associated with +1 nucleosomes of ubiquitous promoters of other animals. 10-bp periodic WW sequences have been observed at +1 nucleosomes in yeast, Drosophila, and zebrafish, but not in mammals (Albert et al. 2007; Mavrich et al. 2008b, 2008a; Tolstorukov et al. 2009; Ioshikhes et al. 2011; Haberle et al. 2014; Wright and Cui 2019). However, whether 10-bp WW periodicity is associated with promoters of particular types has not been investigated.

We first examined TSS sets that represent all genes in Drosophila, zebrafish, mouse, and human (see Methods). As expected, we detected 10-bp WW periodicity signals downstream of Drosophila and zebrafish TSSs, but not human TSSs, and we found that this signal was also not detected in mouse (Fig. 5A). As in C. elegans, we observed that the WW periodicity signals in Drosophila and zebrafish peaked in the 5’ half of +1 nucleosomes (Fig. 5A).

**Figure 5.**
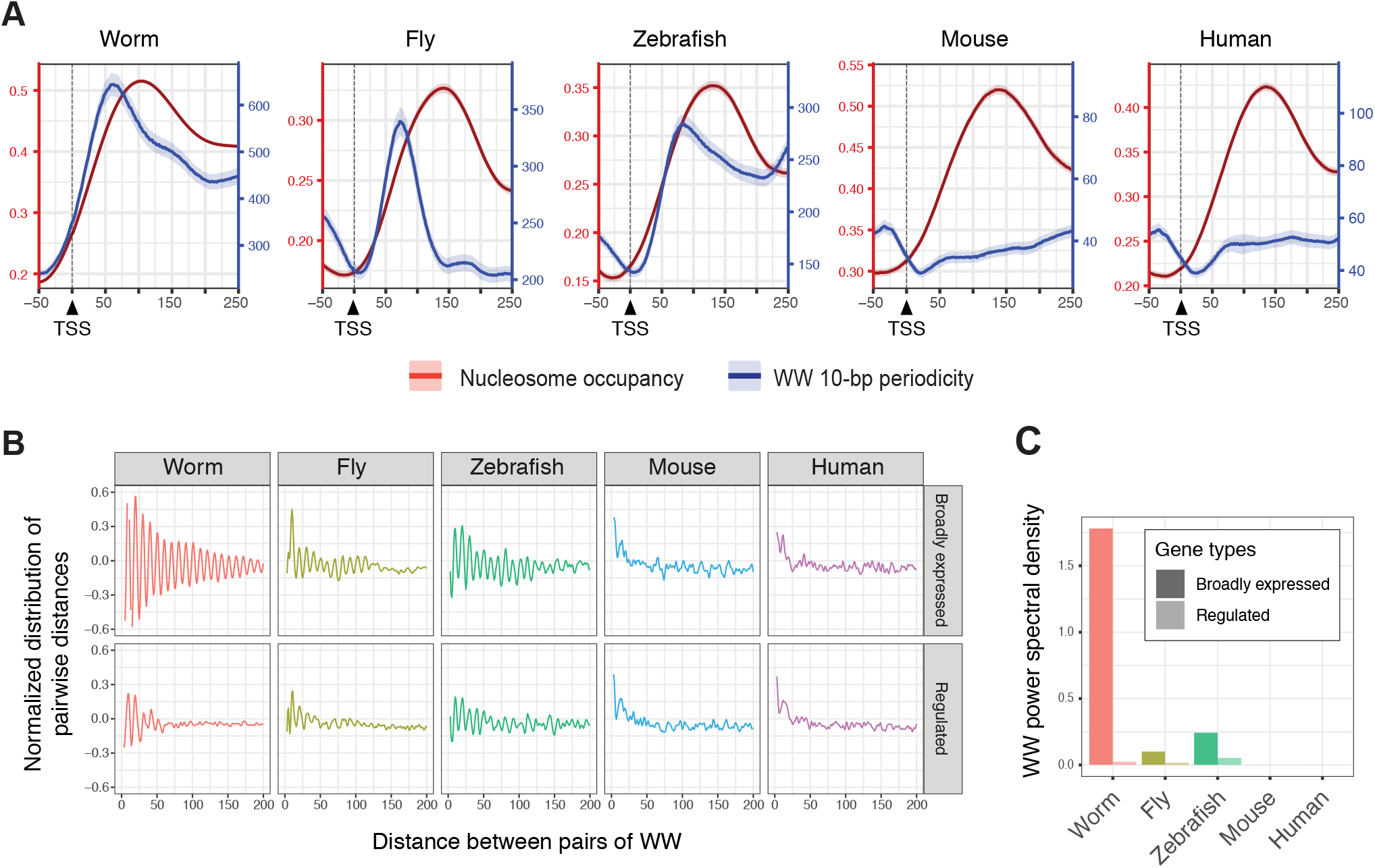
10-bp WW periodicity at ubiquitous promoters is a feature of non-mammalian genomes. (A) Nucleosome occupancy probability scores (red, left axis) and 10-bp WW periodicity (blue, right axis) at worm, fly, zebrafish, mouse and human TSSs. (B) Normalized distribution of pairwise distances between WW dinucleotides found in the sequences from −50 bp to +300 bp relative to TSSs, for genes with broad expression (top row, 20% lowest gene expression cv scores) or regulated expression (bottom row, 20% highest gene expression cv scores) in worm, fly, zebrafish, mice and human. (C) Associated WW power spectral densities values at a 10-bp period.

We then investigated subsets of promoters to ask whether 10-bp WW periodicity signals are associated with ubiquitously active promoters and to compare with signals at promoters with regulated activity. Using the coefficient of variation of gene expression (cv) as a metric, we considered genes in the bottom 20% of cv values to have broad ubiquitous expression and those in the top 20% to have highly regulated expression (e.g., tissue specificity). As found in C. elegans, we observed that promoters of broadly expressed genes in Drosophila and zebrafish have higher 10-bp WW periodicity signals than those of highly regulated genes. In contrast, neither the broadly active nor the regulated groups of mouse and human promoters had detectable WW periodicity signals (Fig. 5B-C). These results suggest that 10-bp WW periodicity signals are a conserved feature of ubiquitously active promoters in non-mammalian animals.

## Discussion

Determining regulatory architectures that drive gene expression patterns is necessary for understanding how the genome encodes development. Through comprehensive analyses of gene expression and chromatin accessibility in five adult *C. elegans* tissues covering ∼90% of cells, we show that most genes have either ubiquitous or tissue-specific expression and we describe extensive differences between their regulatory architectures. The expression of ubiquitous genes involved in basic biological processes as well as that of germline-specific genes is often controlled by single promoters, whereas soma-specific and ubiquitous genes involved in developmental processes tend to have multiple alternative promoters and enhancers. We also found that the majority of regulatory elements have tissue-specific accessibility and we identified differences in sequence composition between promoters active in different tissues.

We found that a strong +1 nucleosome position coinciding with a 10-bp periodic WW signal is a key feature of ubiquitous and germline-specific promoters in *C. elegans*. The association of 10-bp WW periodicity and nucleosome rotational position was first noted by Travers and colleagues in chicken, and is thought to aid nucleosome positioning by conferring sequence-dependent bendability to the DNA polymer (Zhurkin et al. 1979; Trifonov 1980; Drew and Travers 1985). Such periodicity has been observed in nucleosomal sequences in different eukaryotes including *C. elegans* but its specific association with different gene types was unknown (Satchwell et al. 1986; Ioshikhes et al. 1996; Widom 2001; Segal et al. 2006; Johnson et al. 2006; Peckham et al. 2007; Field et al. 2008; Mavrich et al. 2008a, 2008b; Ioshikhes et al. 2011; Struhl and Segal 2013; Forrest et al. 2014; Haberle et al. 2014; Dreos et al. 2016; Pich et al. 2018).

In contrast to ubiquitous and germline-specific promoters, +1 nucleosomes of somatic tissue-specific promoters are not associated with a 10-bp WW periodicity signal, have lower occupancy, and inconsistent position relative to the TSS. Instead, we observed intriguing biases in the enrichment of core motifs at these promoters. TATA boxes are primarily found in hypodermal and intestinal promoters whereas Sp1 motifs are most highly enriched in neuronal promoters. In addition, tissue-specific motifs are present, and these often have peak positions around −50bp relative to the mode TSS.

Structural studies of the Pre-Initiation Complex (PIC) showed that it covers the region from about −45 bp to +20 bp relative to the transcription start site (Louder et al. 2016; Robinson et al. 2016; Schilbach et al. 2017). Interestingly, the 5’ edges of the +1 nucleosomes at *C. elegans* ubiquitous and germline promoters are located ∼ 20 bp downstream of the TSS, which would be at the 3’ edge of the PIC. This supports the model initially proposed in yeast whereby a positioned +1 nucleosome could facilitate PIC complex assembly by interacting with TFIID (Jiang and Pugh 2009) (Fig. 6). At soma-specific promoters, which lack strongly positioned nucleosomes, the binding of core or tissue specific TFs ∼ 45 bp upstream of the TSS might help to locally recruit and/or to position the PIC (Fig. 6). These models are not mutually exclusive and additional mechanisms also contribute to promoter activity.

**Figure 6.**
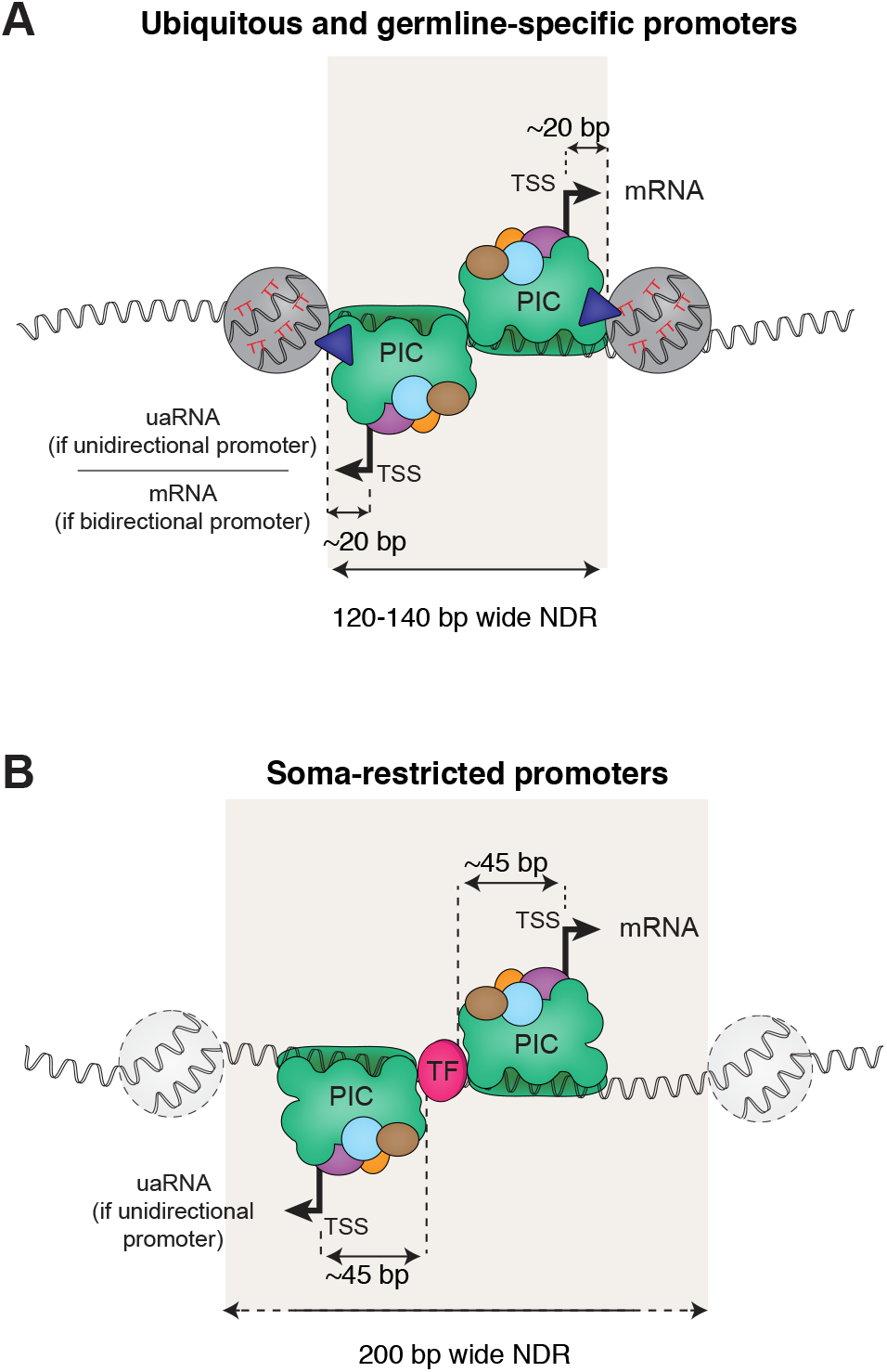
Two models of PIC positioning at promoters. The nucleosome organization and sequences features found in ubiquitous, germline-specific and somatic-tissue-specific promoters suggest that two models of Pre-Initiation Complex recruitment exist. (A) In ubiquitous and germline-specific promoters (*i.e.* germline-active promoters), nucleosomes flank a narrow 120 to 140 bp-wide NDR. Positioning of these nucleosomes is facilitated by the underlying DNA sequence which harbors highly periodic WW (mainly TT) dinucleotides. Thus, the Pre-Initiation Complex (PIC) assembling at the NDR is physically constrained by the +1 nucleosome edge, resulting in transcription initiation ∼20 bp upstream of the +1 nucleosome edge. Many of these promoters lead to bidirectional elongative transcription. Otherwise, upstream-antisense RNA (uaRNA) are transcribed. (B) In soma-restricted promoters, NDRs are wider (> 200 bp) and flanking nucleosomes are weakly positioned and not reproducibly aligned relative to the TSS. Core and transcription factors recruited to the NDR facilitate assembly and positioning of the PIC, resulting in transcription initiation −45 to −50 bp downstream.

Similar to *C. elegans*, we observed that a 10-bp WW periodicity signal is also associated with promoter +1 nucleosomes of broadly expressed genes in zebrafish and Drosophila. This is consistent with a previously described enrichment of 10-bp periodicity in AA and TT dinucleotides downstream of zygotic TSSs in zebrafish (Haberle et al. 2014). A weak genome-wide AA/TT periodicity was previously noted in Drosophila but not associated with any gene feature (Mavrich et al. 2008b). In contrast, the periodic WW signal is not detected at promoters of broadly expressed genes in mouse and human, despite their having well positioned +1 nucleosomes. This is consistent with reports showing a low 10-bp WW periodicity in mammal genomes, either around TSSs (Tolstorukov et al. 2009; Wright and Cui 2019) or genome-wide (Pich et al. 2018). Multiple factors have been shown to contribute to nucleosome positioning in eukaryotes, including intrinsic DNA sequence, chromatin remodelers, DNA binding proteins, and RNA polymerase machinery (Jiang and Pugh 2009; Struhl and Segal 2013). We suggest that 10-bp WW periodicity is an ancient conserved signal that contributes to +1 nucleosome positioning at ubiquitously active promoters of non-mammalian eukaryotes, especially those of genes with basal cell functions, whereas nucleosome positioning in mammals may rely on other mechanisms (Struhl and Segal 2013).

In addition to illuminating understanding of regulatory architectures, we provide extensive datasets and annotation of gene expression and accessible chromatin across adult tissues, available at the *C. elegans* regulatory atlas (RegAtlas, https://ahringerlab.com). These data and tools will be key resources that facilitate future studies of *C. elegans* gene expression regulation by the scientific community.

## Material and methods

### Nuclear sorting

Young adult hermaphrodites were obtained by growing synchronized starved L1 larvae at 25 C in standard S-basal medium with HB101 bacteria for 40-42h. After sucrose flotation and washing in M9 buffer, worms were frozen into “popcorn” by dripping concentrated slurry into liquid nitrogen. Nuclei were isolated as previously detailed (Jänes et al. 2018), with minor modifications. ∼ 20,000 to 200,000 frozen young adult worms were broken by smashing using a Biopulverizer then the frozen powder was thawed in 8 ml Egg buffer (25 mM HEPES pH 7.3, 118 mM NaCl, 48 mM KCl, 2 mM CaCl2, 2 mM MgCl2). Broken worms were pelleted by spinning at 800 g for 3 min then resuspended in 8 ml of Buffer A (0.3 M sucrose, 10 mM Tris pH 7.5, 10 mM MgCl2, 1 mM DTT, 0.5 mM spermidine 0.15 mM spermine, protease inhibitors (Roche complete, EDTA free) and 0.025 % IGEPAL CA-630). The sample was dounced (two strokes) in a 14-ml stainless steel tissue grinder (VWR) then spun at 100 g for 6 min to pellet remaining worm fragments. The supernatant was kept (nuclei batch 1) and the pellet resuspended in a further 7 ml of Buffer A and dounced for 30 strokes. This was spun at 100 g for 6 min to pellet debris and the supernatant was kept (nuclei batch 2). The first fraction was enriched for germline nuclei while the second fraction was enriched for somatic nuclei. Nuclei quality was assessed by microscopy.

Following isolation, nuclei were immunostained by adding phycoerythrin-coupled anti-GFP antibody (Biolegend # 338003) at 1:200 in 7 ml of buffer A, and 280 units of murine RNAse inhibitor (M0314S) were added to protect RNA from being degraded. Nuclei were kept slowly rotating at 4 C in the dark for 1 to 16 hours. Debris was removed by spinning at 100 g for 6 min at 4 C then nuclei were pelleted (2000 g for 20 min at 4 C), washed in 6 ml of buffer A, and resuspended in buffer A containing 80 U/ml murine RNAse inhibitor at a concentration of ∼ 10-15 million nuclei / ml. Finally, nuclei were filtered on 30 µm mesh (CellTrics 04-0042-2316) and stained with 0.025 µg/ml DAPI. Nuclei quality was assessed immediately before sorting by microscopy.

Nuclear sorting was performed at 4 C using a Sony SH800Z sorter fitted with a 100 µm sorting chip and auto-calibrated. Nuclei were gated using the DAPI signal and PE-positive nuclei were gated using PE-H / BSC-A signal. DAPI gating depended on which nuclei were being sorted (*e.g.* intestine nuclei are 32N). A recording speed > 15,000 nuclei per second ensured a sorting efficiency higher than 80 %. Nuclei were sorted into 15 ml Falcon tubes containing 500 µl of buffer A with 800U/ml murine RNAse inhibitor. Nuclei were sorted in batches of one million and then processed for downstream applications. The purity and integrity of each batch of nuclei was assessed by recording an aliquot of sorted nuclei in a second pass in the sorter and by microscopy. All sorted samples used in this study had a purity higher than 95%.

### ATAC-seq

One million sorted nuclei were pelleted (2000 g for 20 min at 4 C) and resuspended in 1X Tn5 Buffer (10mM Tris pH 8, 5mM MgCl2, 10% DMF) at a final concentration of ∼ 500,000 nuclei / ml. 2.5 ul of Tn5 (Illumina FC-121-1030) were added to 47.5 ul (∼ 25,000 nuclei) of the suspension. ATAC-seq was then performed as previously described (Jänes et al. 2018). ATAC-seq libraries were generated from two biological replicates for each tissue, and were sequenced in both single-end and paired-end modes. Single ATAC-seq libraries were made for L1 and L3 muscle (SE-sequenced) and L3 germline (PE-sequenced). PGC-specific ATAC-seq data at the L1 stage was obtained from (Lee et al. 2017).

### RNA-seq

RNA was extracted from one million sorted and washed nuclei using standard procedure (Jänes et al. 2018). A minimum of 20 ng of total nuclear RNA were used to make long nuclear RNA-seq libraries. Long nuclear RNA (>200 nt) was isolated using Zymo Clean and Concentrate columns (#R1013), rRNA was removed using the Ribo-Zero rRNA removal kit (MRZH11124), and stranded libraries were prepared with the NEBNext Ultra Directional RNA Library Prep Kit (#E7420S). Long nuclear RNA-seq libraries were generated from two biological replicates for each tissue and were sequenced in paired-end mode. We observed that all tissue-specific libraries had contamination of whole animal cytoplasmic RNA presumably bound to nuclei during initial nuclei released. This contamination caused all samples to have noticeable background for abundant mRNAs (*e.g.*, muscle myosin *unc-54*).

### Data processing

Data was processed as described in (Jänes et al. 2018; Chen et al. 2013) and aligned to WBcel235/ce11 genome. Further details are given in the Supplemental methods. To assess the reproducibility of biological replicate datasets, we used site accessibility or gene expression values to compute pairwise Euclidean distances between each dataset and pairwise Pearson correlation scores. ATAC-seq and RNA-seq biological replicates showed high concordance (Supplemental Fig. S2D).

### Classification of accessible sites

ATAC-seq reads were assigned to the 47,514 annotated accessible sites (150bp width) using the summarizeOverlaps() function from the GenomicAlignments package. Estimation of accessibility fold-changes (FC) and adjusted p-values were computed between all pairs of tissues using the DESeq2 package (Love et al. 2014). A site was considered significantly differentially accessible (DA) between two tissues if there was a fold-change > 3 and an adjusted p-value < 0.01. A fold change of 3 between consecutive tissues was used as threshold to determine the tissue specificity of accessible sites. Classification details are provided in the Supplemental Material.

### Classification of genes

Long nuclear RNA-seq stranded fragments were assigned to *C. elegans* gene annotations (WBCel235, release 92) using the featureCounts program with “-t gene -s 2 -Q 10 -p” options. Estimation of expression fold-changes (FC) and adjusted p-values were computed between pairs of tissues using the DESeq2 package. A gene was considered significantly differentially expressed (DE) between two tissues if there was a fold-change > 3 and an adjusted p-value < 0.01.

In each sample, gene expression was calculated as Transcripts Per Million (TPM) values. TPMs of biological duplicates were then averaged to obtain a single gene expression value for each tissue. The rules used to classify accessible sites (detailed in the Supplemental methods) were also used to classify genes, with a detection threshold of 5 TPM. A small number of germline-specific genes (151) with maximal expression in L4 (Jänes et al. 2018) were classified as sperm-specific and not included in this study.

### GO analysis

GO enrichment analyses were performed using the gProfileR 0.6.7 package (Reimand et al. 2007), filtering for redundant GO terms using the hier_filtering = moderate option. To compare GO enrichment across several groups, the clusterProfiler 3.10.1 package (Yu et al. 2012) was used, filtering for redundant terms using REVIGO. Only GO terms with Bonferroni-adjusted p-values lower than 0.05 were kept.

### ATAC-seq fragment density plots

ATAC-seq fragment density plots, also known as V-plots (Henikoff et al. 2011), were generated using the VplotR 0.4.0 package (https://github.com/js2264/VplotR). Flanking nucleosome enrichment scores were calculated from the V-plots as illustrated in Supplemental Fig. S5C.

### Nucleosome occupancy tracks and putative +1 nucleosome mapping

Processed bam files from paired-end ATAC-seq duplicates of each tissue or from whole organism young adults (Jänes et al. 2018) were merged. For each class of promoter (germline, neuron, muscle, hypodermis, intestine and ubiquitous promoters), the nucleoATAC python package (Schep et al. 2015) was used to compute the probability of nucleosome occupancy from −1kb to + 1kb from promoter centers in each tissue (germline, neuron, muscle, hypodermis, intestine and whole organism).

Putative +1 and −1 nucleosome positions were determined for each set of tissue-specific promoters using the corresponding tissue-specific nucleosome occupancy probability track and for ubiquitous promoters using whole organism nucleosome occupancy probability track (Jänes et al. 2018). We assigned the center of the putative +1 nucleosome to the local maximum of the nucleosome occupancy probability within 200 bp downstream from the forward TSS mode. Similarly, the center of the −1 nucleosome summit was assigned to the local maximum of the occupancy probability within 200 bp upstream of the reverse TSS mode. Only coding promoters with experimentally determined forward and reverse TSSs were considered.

### Motif identification and enrichment analyses

Motifs enriched in different sets of promoters (−75 bp to +105 bp from promoter centers) were identified using MEME in stranded mode and a 0-order background model (-markov_order 0). MEME mode was set to ‘Any Number of Repetitions’ (-mod anr) and motif widths were restricted to 6 to 25 bp. The five motifs found most enriched (with an E-value threshold of 0.05) were retrieved. Unstranded motifs (found twice as complementary sequences, since MEME was run in stranded mode) were manually combined. PWMs for the Initiator (Inr) and the TATA motif were obtained from (Jin et al. 2006). Motif mapping to promoters was performed in R using the Biostrings 2.50.2 package, the GenomicRanges 1.34.0 package and the TFBSTools 1.20.0 package with a relScore threshold set to 0.8.

### Dinucleotide periodicity

To estimate dinucleotide periodicity in sets of sequences (*e.g.* −50 to +300 bp sequences around ubiquitous, germline or somatic-tissue-specific TSSs in Fig. 4B, or −50 to +300 bp sequences around TSSs from different organisms in Fig. 5A), the getPeriodicity() function from the periodicDNA 0.2.0 package was used with default parameters. Briefly, the distribution of distances between all possible pairs of dinucleotides in the set of sequences was computed and corrected for distance decay, smoothed by a moving average window of 3 and power spectral densities were retrieved by applying a Fast Fourier Transform to the normalized distribution.

To generate 10-bp dinucleotide periodicity score tracks, the generatePeriodicityTrack() function from the periodicDNA 0.2.0 package (https://github.com/js2264/periodicDNA) was used with default parameters. Briefly, a running 10-bp dinucleotide periodicity score was calculated by applying a Fast Fourier Transform (stats 3.5.2 package) on the distribution of distances between pairs of dinucleotides (*e.g.* WW……WW) found in 100-bp long sequences (2-bp increments).

### Phasing of nucleosomal sequences

To observe the 10-bp periodic occurrence of a dinucleotide in putative +1 nucleosomes, sequences (400 bp centered at the nucleosome dyads) were first clustered by k-means based on the dinucleotide occurrences in each sequence, then the clusters were rephased within a −/+5 bp range using the lag value estimated by the ccf() function from the stats 3.5.2 package.

### Sets of annotations in fly, fish, mouse and human

In worms, experimentally annotated TSSs were used (Jänes et al. 2018). In fly and zebrafish (respectively dm6 and danRer10 genome versions), TSSs were assigned to the first base of the genes using TxDb.Dmelanogaster.UCSC.dm6.ensGene 3.4.4 and TxDb.Drerio.UCSC.danRer10.refGene 3.4.4 gene models with the GenomicFeatures 1.34.7 package in R. In mouse and human, FANTOM CAGE datasets (Lizio et al. 2015) were used to retrieve the dominant TSS closest to the gene annotation.

Coefficient of variation of gene expression (CV) values were retrieved from (Gerstein et al. 2014) for worm, fly and human or computed using gene expression datasets from (Pervouchine et al. 2015) for mouse and (White et al. 2017) for zebrafish. Genes with the 20% lowest CVs were considered broadly expressed and those with the 20% highest CVs were considered regulated.

### Nucleosome occupancy in metafly, fish, mouse and human

Nucleosome occupancy tracks were generated as described for worms using nucleoATAC with the following ATAC-seq datasets: SRR6171265 (Haines and Eisen 2018) in fly, SRR5398228 (Quillien et al. 2017) in zebrafish, SRR5470874 (Benchetrit et al. 2019) in mouse and SRR891268 (Buenrostro et al. 2013) in human.

### Other visualization tools

Figures were generated in R 3.5.2, using either base or ggplot2 3.1.1 plotting functions. Genome browser screenshots were obtained from IGV 2.4.8. Genome tracks in the bigWig format were imported in R using the rtracklayer 1.42.2 package.

### Data and software availability

Processed data and all annotations are available and can be either dynamically explored or anonymously downloaded at https://ahringerlab.com/. Key analysis concepts developed for this study have been integrated into two R packages available at https://github.com/js2264/VplotR/ and https://github.com/js2264/periodicDNA/.

Raw and processed sequencing datasets generated in this paper are available at GEO under the accession number GSE141213.

## Acknowledgments

We thank all the members of the Ahringer lab and particularly F. Carelli and A. Frapporti for fruitful discussions and comments on the manuscript. We thank A. Appert and K. Harnish for support in sequencing. The work was supported by a Wellcome Trust Senior Research Fellowship to J.A. (101863) and a Medical Research Council DTP studentship to J.S.. We also acknowledge core support from the Wellcome Trust (092096) and Cancer Research UK (C6946/A14492).

## Author contributions

J.S. and J.A. conceived and designed the studies. JS performed experiments unless otherwise noted. J.S., M.C. and C.C. generated the reporter strains. J.S. and Y.D. prepared RNA-seq libraries. J.S. and J.J. processed the data. J.S. analyzed the data. J.S. and J.A. wrote the manuscript. J.A. supervised the study and provided funding.

## Supplemental methods

### Transgenic strains

*C. elegans* strains were maintained using standard procedures at 25°C and fed OP50 *E. coli*. Targeting of the GFP to the nuclear envelope was achieved in two different ways: 1) by fusing a StrepTag (WSHPQFEK) to the N-terminal extremity of GFP (from pPD95.02, Fire lab Vector Kit) and UNC-83 (aa 1-290) to its C-terminal extremity, or 2) fusing the full-length NPP-9 coding sequence to the C-terminal extremity of GFP. The first approach was used to target GFP to the nuclear envelope in germline, muscle, hypodermis and intestine cells. The second approach was used to target GFP to the nuclear envelope in neurons. The promoter used to express the reporter in individual tissues are the *mex-5* promoter (for Germline expression, chrIV:13,353,242-13,353,729), the *egl-21* promoter (for Neuron expression, chrIV:10,481,768-10,481,932), the *myo-3* promoter (from Muscle expression, chrV:12,234,302-12,236,686), the *dpy-7* promoter (for Hypodermis expression, chrX:7,537,794-7,538,688) and the *npa-1* promoter (for Intestinal expression, chrV:7,075,526-7,075,947) (coordinates are in ce11). Three-way Gateway cloning was used to clone each tissue-specific promoter (in slot 1) upstream of the reporter coding sequence (in slot 2). *tbb-2*-3’ UTR was used in slot 3 (Merritt et al., 2008). The destination vector was pCFJ150 (Frøkjaer-Jensen et al., 2008). Reporter constructs were integrated in a single copy at the ttTi5605 Mos1 site located on chrII (Frøkjaer-Jensen et al., 2008).

### Data processing

Reads were trimmed using fastx_trimmer 0.0.14 and aligned to the reference genome WBcel235/ce11 obtained from Ensembl release 92 (ftp://ftp.ensembl.org/pub/release-92/) using bwa-backtrack 0.7.17-r1188 (Li & Durbin, 2009) in single-end (ATAC-seq) or paired-end mode (ATAC-seq, long nuclear RNA-seq). Low-quality (q < 10), mitochondrial and modENCODE-blacklisted (Consortium, 2013) reads were discarded.

Normalized genome-wide accessibility tracks were computed with MACS2 (Feng et al., 2012) using parameters --format BAM --bdg --SPMR --gsize ce --nolambda --nomodel --extsize 150 --shift -75 --keep-dup all and the bedGraphToBigWig utility (Kent et al., 2010). ATAC-seq was also sequenced in paired-end mode; paired-end data were used for nucleosome occupancy and V-plots analyses (described below).

Long nuclear RNA-seq data were processed essentially as in (Chen et al., 2013). Following alignment and filtering, fragments-per-million-normalized strand-specific coverage tracks were computed by transforming the bam file into a bedGraph file using the genomeCoverageBed v2.26.0 utility (Quinlan & Hall, 2010) with the parameters -bg -pc -scale 10e6/${NBFRAGS} -strand ${STRAND} (where ${NBFRAGS} is the number of mapped fragments and ${STRAND} is + or −). Gene annotations used throughout this study are WBcel235/ce11 obtained from Ensembl release 92 (ftp://ftp.ensembl.org/pub/release-92/).

### Annotation of new accessible sites

In a previous study, we identified 42,245 accessible sites across development and aging and annotated them into functional classes (coding promoters, non-coding promoters, unassigned promoters, putative enhancers, inactive elements) based on nuclear RNA seq patterns (Jänes et al., 2018). This annotation pipeline was run using the previously generated data together with the newly generated adult tissue-specific ATAC-seq and RNA-seq generated in this study. This resulted in the detection and annotation of 5,269 new accessible sites, bringing the total sites to 47,514. Supplemental Table S2 provides annotation of the new elements and updated annotations of the elements identified in (Jänes et al., 2018).

### Classification of accessible sites

In each sample, accessibility at each site was calculated as Reads Per Million (RPM) values. RPMs of biological replicates were averaged to obtain a single accessibility score for each site in each tissue. Sites with accessibility lower than 8 RPM in every tissue were not further studied.

The tissue specificity of accessible sites was determined according to the following successive rules:

- *Restricted to a single tissue*: sites (i) significantly DA between the first and the second most accessible tissues and (ii) not significantly DA between the second and the third most accessible tissues.
- *Restricted to two tissues:* sites (i) significantly DA between the second and the third most accessible tissues and (ii) not significantly DA between the third and the fourth most accessible tissues.
- *Restricted to three tissues:* sites (i) significantly DA between the third and the fourth most accessible tissues and (ii) not significantly DA between the fourth and the fifth most accessible tissues.
- *Restricted to four tissues:* sites significantly DA between the fourth and the fifth most accessible tissues.
- *Ubiquitous-biased:* sites (i) significantly DA between any other pair of tissues (*e.g.* first and fourth most accessible tissue) and (ii) detected across all tissues (RPM > 8 in all replicates).
- *Ubiquitous-uniform* (also referred to as simply “uniform”): sites (i) not significantly DA between any pair of tissues and (ii) detected across all tissues (RPM > 8 in all replicates).
- *Unclassified:* sites with accessibility < 8 RPM in some tissues and not significantly DA could not be confidently classified.

### Comparison with other datasets

Tissue-specific gene expression values from nuclear RNA-seq of sorted young adult nuclei were compared to those obtained by single-cell RNA-seq in L2 (Cao et al., 2017) by computing pairwise Euclidean distances between each dataset.

Our gene expression classes were compared to those derived from single-cell RNA-seq in L2 (Cao et al., 2017). There, genes were considered enriched in a given tissue if the expression fold-change between this tissue and the tissue with the second highest expression was higher than 5. Genes were considered detected if their expression was higher than 5 TPM in at least one tissue, and ubiquitous if (i) their expression was higher than 5 TPM across all tissues and (ii) they were not enriched in any tissue. Our gene expression classes were also compared to those obtained by tissue-specific cell sorting and RNA-seq in young adult somatic tissues (muscle, neurons, hypodermis and intestine) (Kaletsky et al., 2018).

## Supplemental figures and associated legends

**Supplemental Figure S1.**
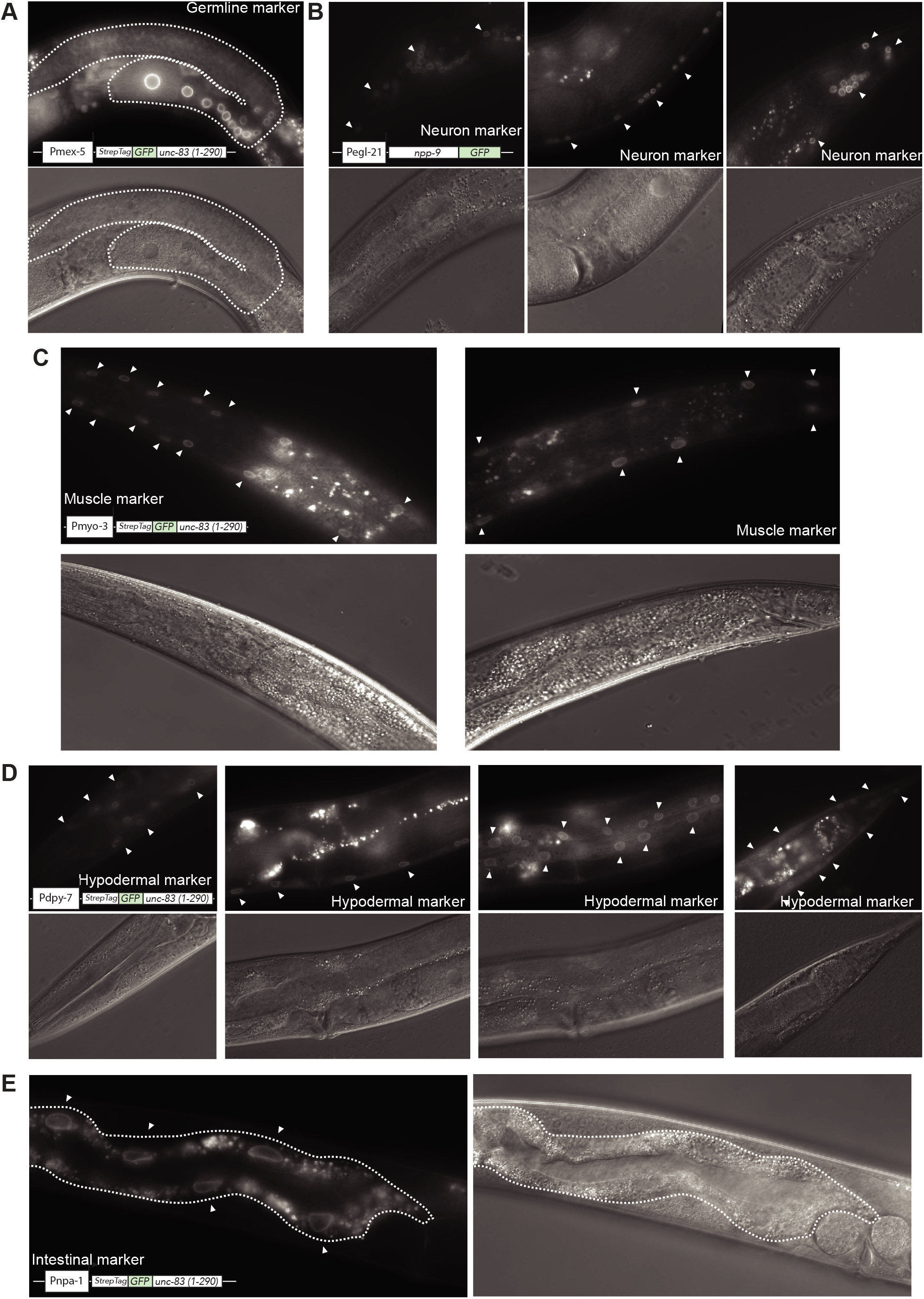
Reporter strains created for this study. Reporter strains labelling nuclear envelope of germline nuclei (A), neuronal nuclei (photos of head neurons, ventral nerve cord and tail neurons) (B), muscle nuclei (photos of anterior and posterior sides) (C), hypodermis nuclei (photos of head, ventral hypodermal ridge, seam and tail) (D) and intestine nuclei (photos of anterior intestine) (E). For each reporter, the construct used to drive expression of the marker is depicted. DIC images are also shown for reference (bottom).

**Supplemental Figure S2.**
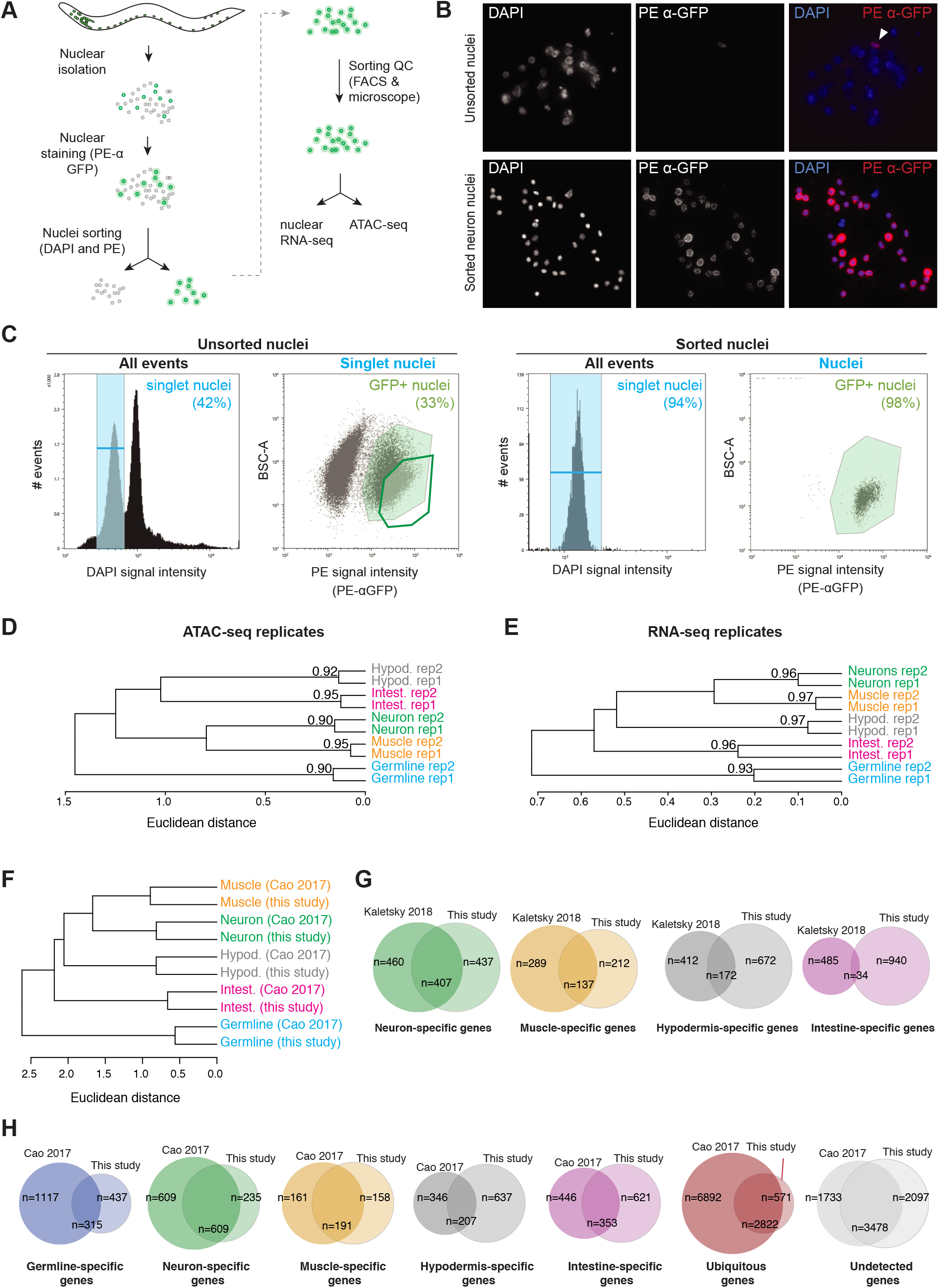
Sorting strategy and datasets quality control. (A) Detailed procedure used to isolate tissue-specific nuclei. (B) Nuclei from neuronal reporter strain (Pegl-21::npp-9::GFP::tbb2-3’UTR) immuno-stained with a PE α-GFP antibody, before (top) and after (bottom) nuclei sorting. The arrow points to a single PE+ nucleus. (C) Left: gating strategy to isolate PE+ (*i.e.* GFP+) nuclei from a nuclear preparation. Single nuclei are gated (shaded blue area) and GFP+ nuclei (green shaded area) are readily separated from GFP-nuclei. Here, the gate used to sort GFP+ nuclei is the thick-lined green gate (no shading). Right: flow cytometry recording of sorted nuclei to estimate the purity of GFP+ nuclei. (D-F) Euclidean distances and Pearson correlation scores between ATAC-seq biological duplicates (D), RNA-seq biological duplicates (E), and between RNA-seq (this study) and single-cell RNA-seq from the L2 stage (Cao et al., 2017) (F). (G-H) Intersection between gene expression annotations (this study) and those from RNA-seq in YA (Kaletsky et al., 2018) or single-cell RNA-seq in L2 (Cao et al., 2017).

**Supplemental Figure S3.**
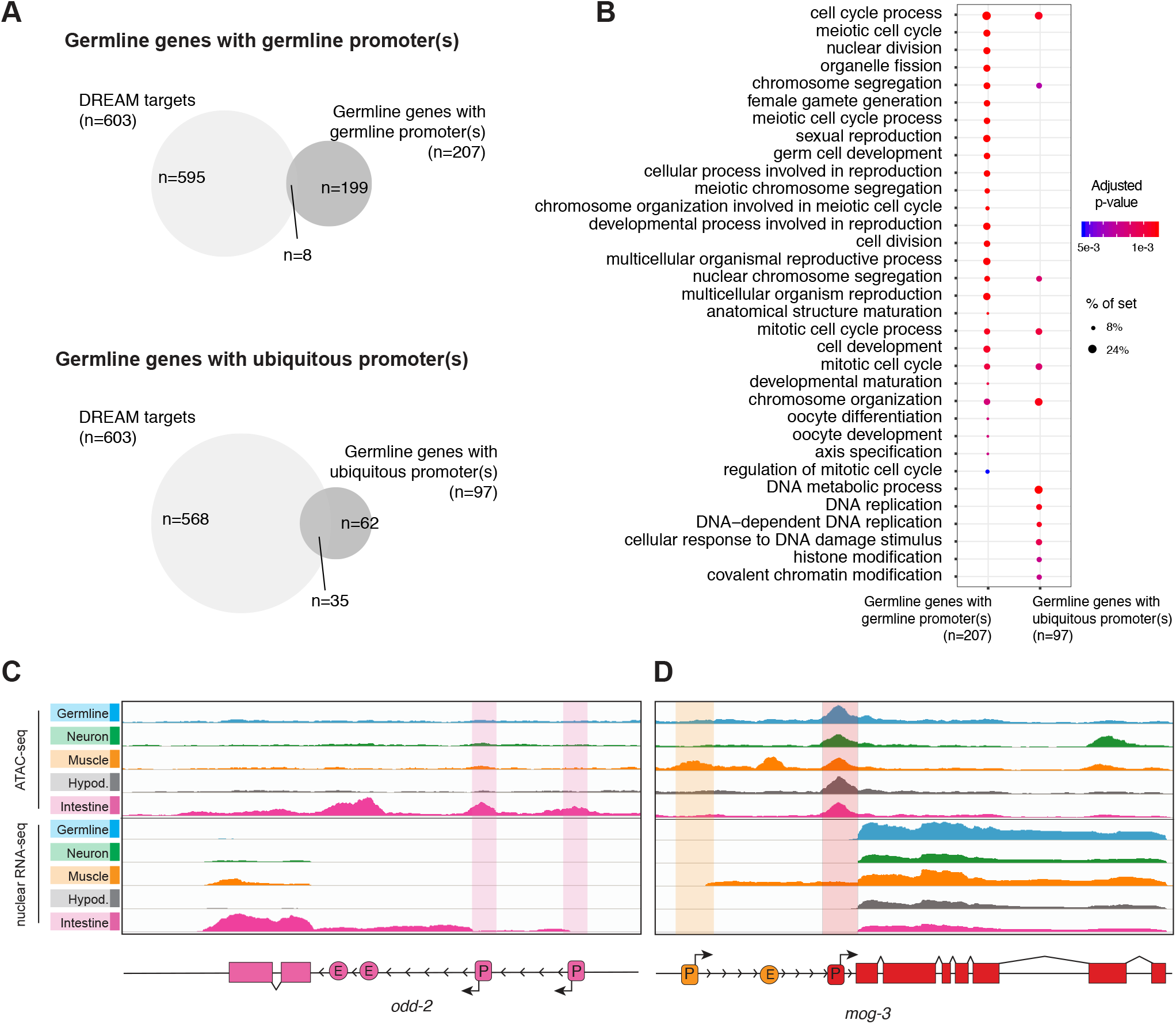
Promoter classes associated with different gene types. (A) Intersection of DREAM targets defined in (Latorre et al., 2015) with germline genes with only germline-specific promoter(s) (top) or only ubiquitous promoter(s) (bottom). (B) GO terms enriched in germline genes with only germline-specific or only ubiquitous promoter(s). (C) Example of a tissue-specific gene with multiple tissue-specific promoters (here odd-2, an intestine gene with two intestine-specific promoters). (D) Example of a ubiquitous gene with a ubiquitous promoter and a tissue-specific promoter (here mog-3, with one ubiquitous and one muscle-specific promoter).

**Supplemental Figure S4.**
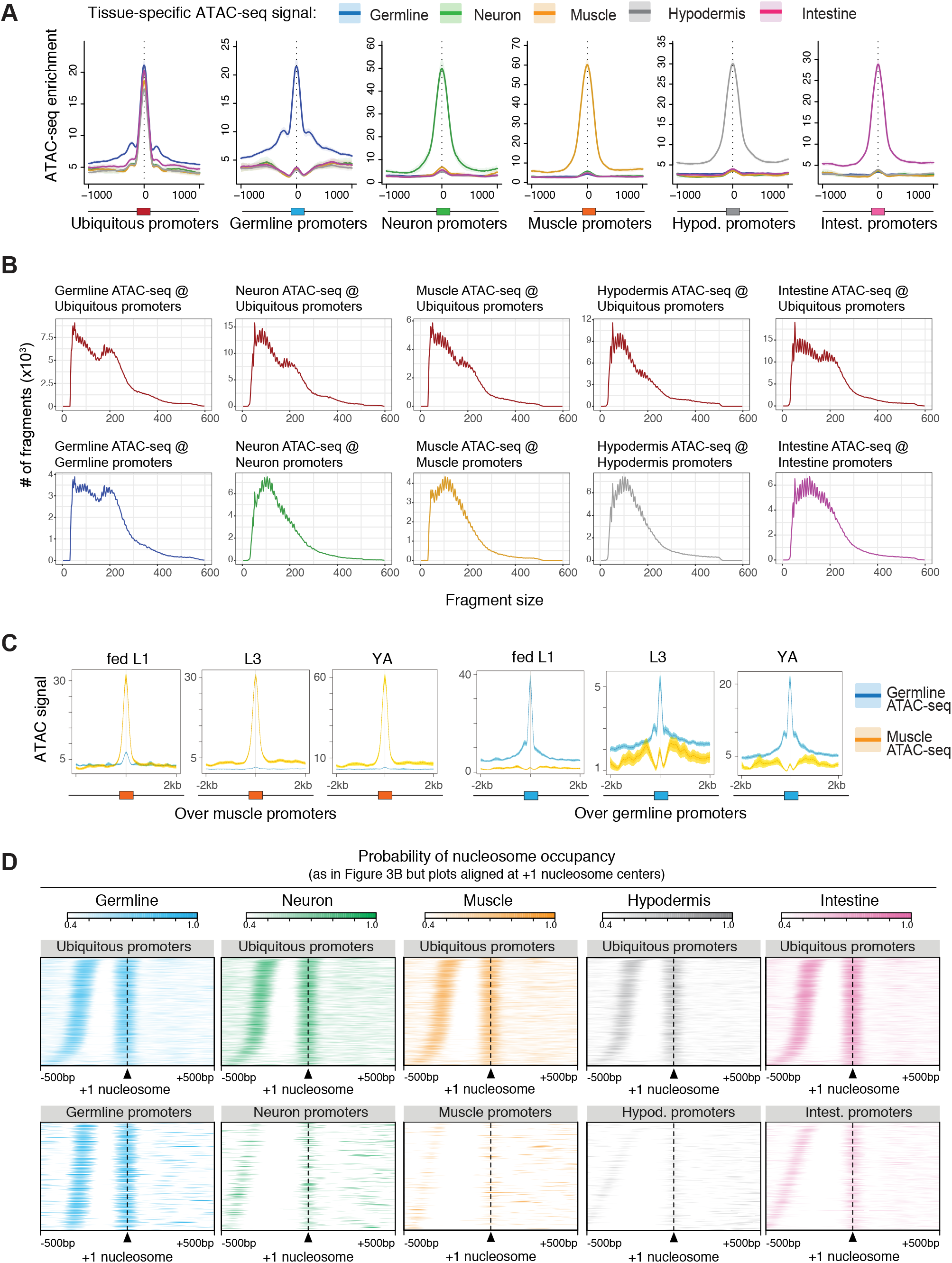
Nucleosome signatures at different types of promoters. (A) Metaplots of tissue-specific ATAC-seq tracks over different classes of promoters. (B) Size distribution of ATAC-seq fragments from different tissue-specific datasets, mapping over ubiquitous or tissue-specific promoters. (C) Metaplots of germline and muscle-specific ATAC-seq tracks obtained at multiple developmental stages (L1, L3 and young adult) over germline or muscle-specific promoters. (D) Same figure as in Figure 3B, but with nucleosome occupancy signals centered at +1 nucleosome summits rather than at TSSs. Rows are ordered by NDR widths.

**Supplemental Figure S5.**
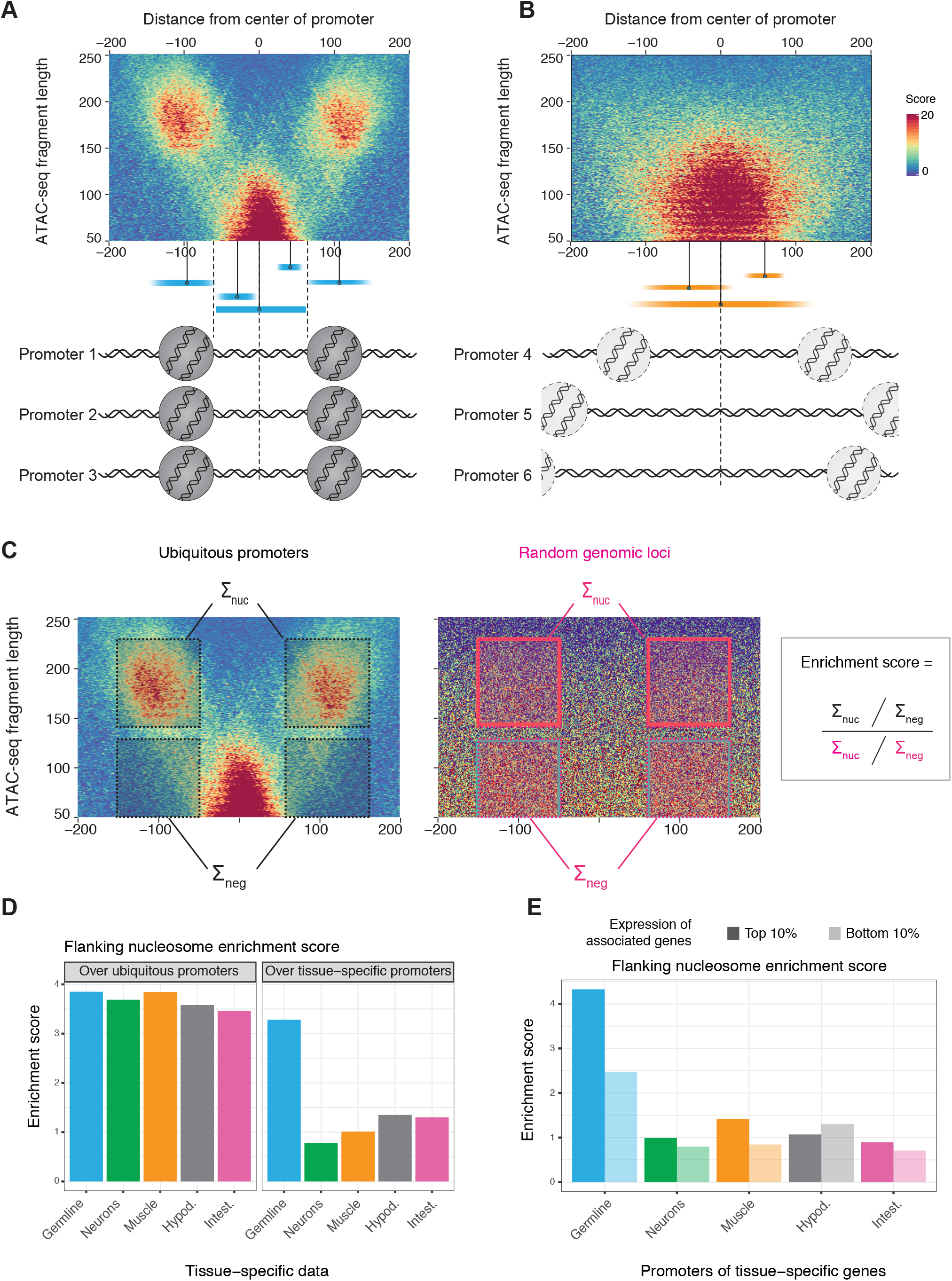
Fragment density plots and flanking nucleosome enrichment scores. (A-B) Interpretation of two ATAC-seq fragment density plots shown in Figure 3. The dense cluster of short fragments at the promoter centers represents the nucleosome-depleted region (NDR) while the two dense clusters of longer fragments located −100 and +100 bp away from the promoter centers are indicative of aligned −1/+1 flanking nucleosomes. (C) Methodology to compute flanking nucleosomes enrichment scores from ATAC-seq fragment density plots. (D) Flanking nucleosomes enrichment scores calculated using different tissue-specific ATAC-seq datasets, at ubiquitous or tissue-specific promoters. (E) Flanking nucleosomes enrichment scores at promoters associated with either the 10% most expressed tissue-specific genes (dark bars) or the bottom 10% least expressed tissue-specific genes (light bars). Note that promoters of lowly expressed germline-specific genes have an enriched +1 nucleosome whereas promoters of soma-restricted highly expressed genes do not.

**Supplemental Figure S6.**
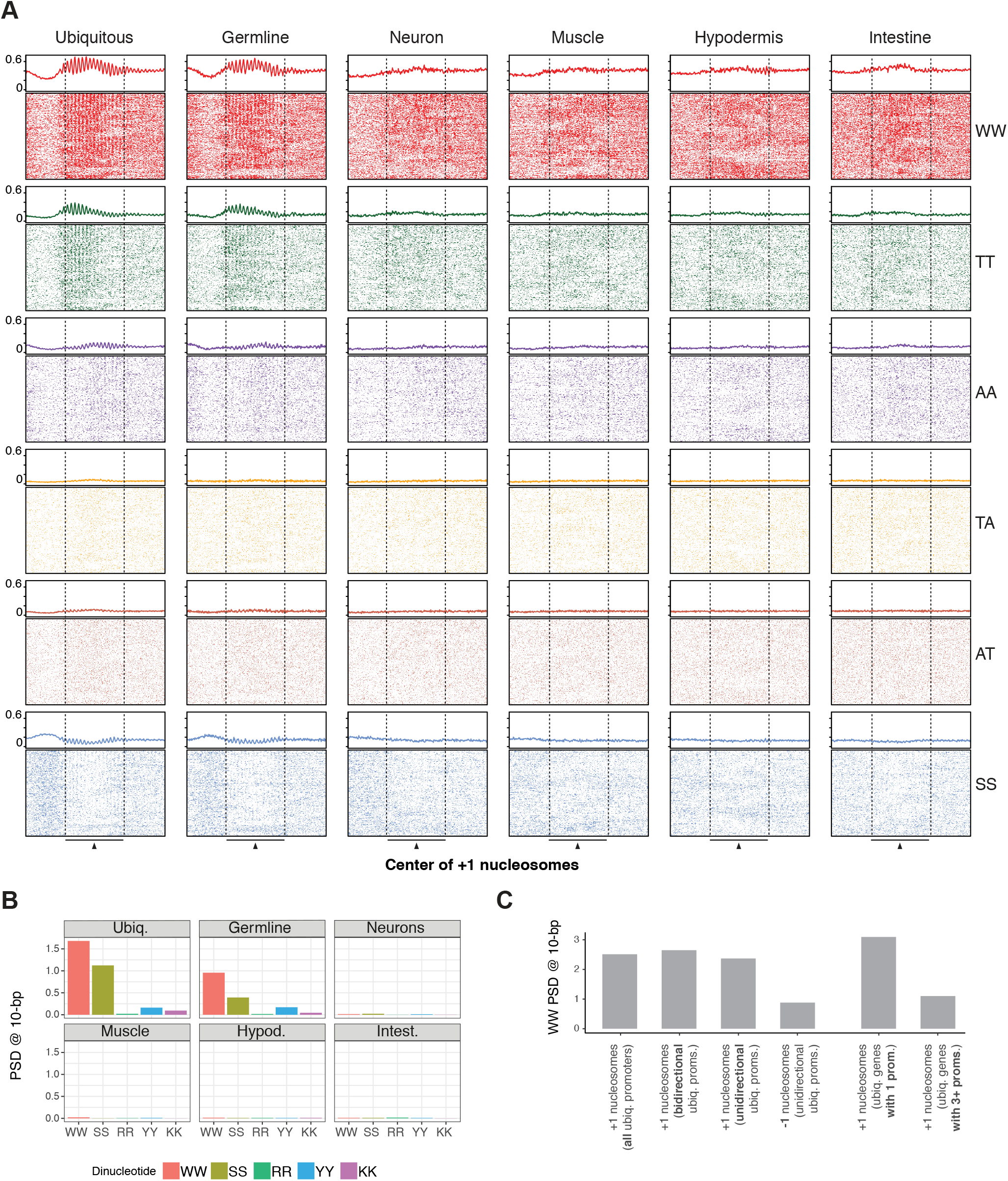
10-bp dinucleotide periodicities at different classes of promoters. (A) WW, TT, AA, TA, AT and SS dinucleotide occurrences observed at +1 nucleosomes of ubiquitous or tissue-specific promoters (400 bp window centered at nucleosome dyads). Rows were shifted up to 5 bp to highlight the phased 10-bp periodic patterns. Summed dinucleotide occurrences are represented on top of each heatmap by a line plot. (B) WW power spectrum density (PSD) values at a 10-bp period for different dinucleotides in +1 nucleosome sequences of ubiquitous and tissue-specific promoters. (C) WW PSD values at a 10-bp period at +1 nucleosomes of different sets of ubiquitous promoters and at −1 nucleosomes of unidirectional ubiquitous promoters.

**Supplemental Figure S7.**
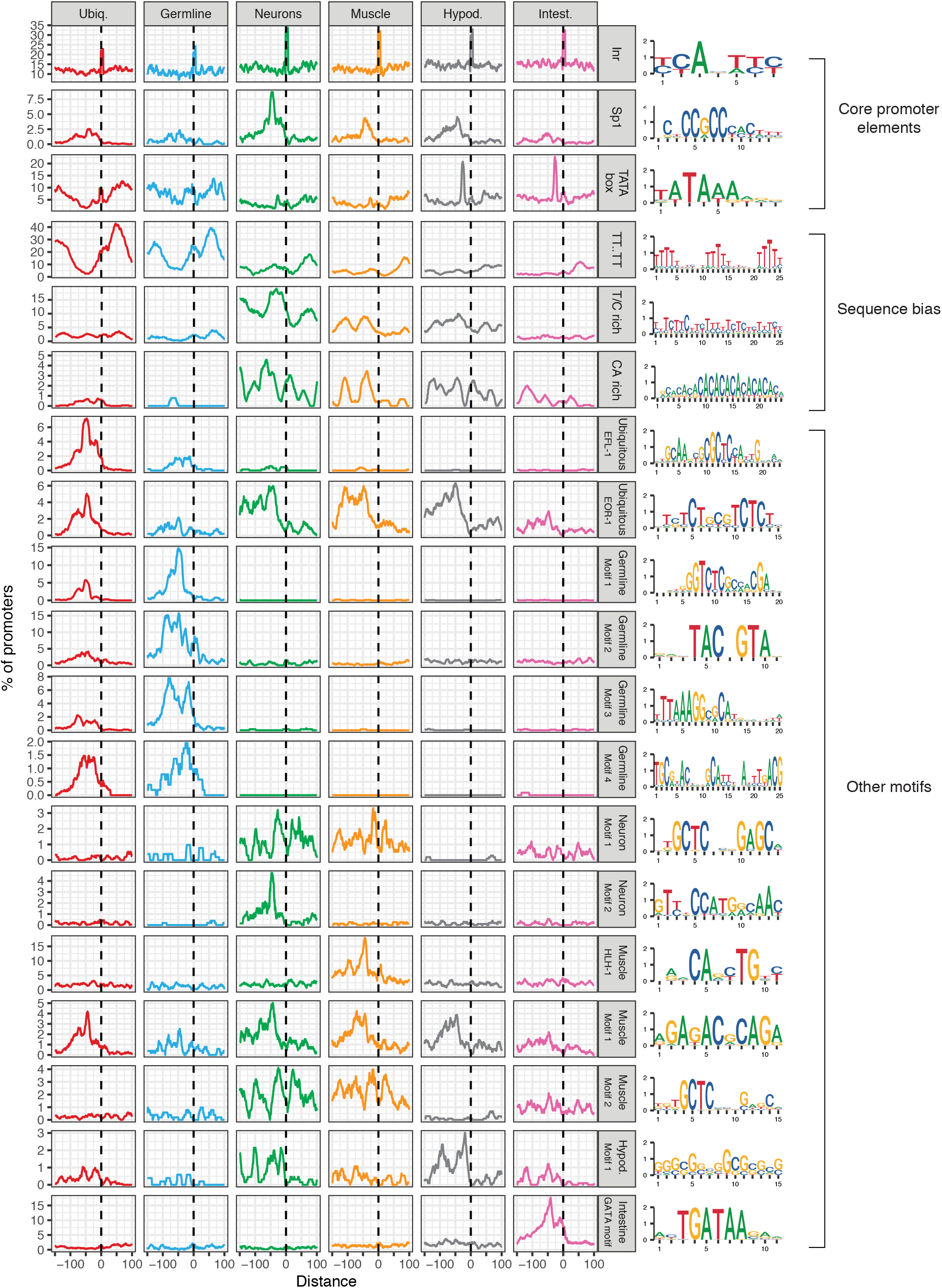
Location of motifs relative to ubiquitous or tissue-specific TSSs. Motif PWMs are displayed on the right. Only promoters with experimentally defined TSSs were considered.

**Supplemental Figure S8.**
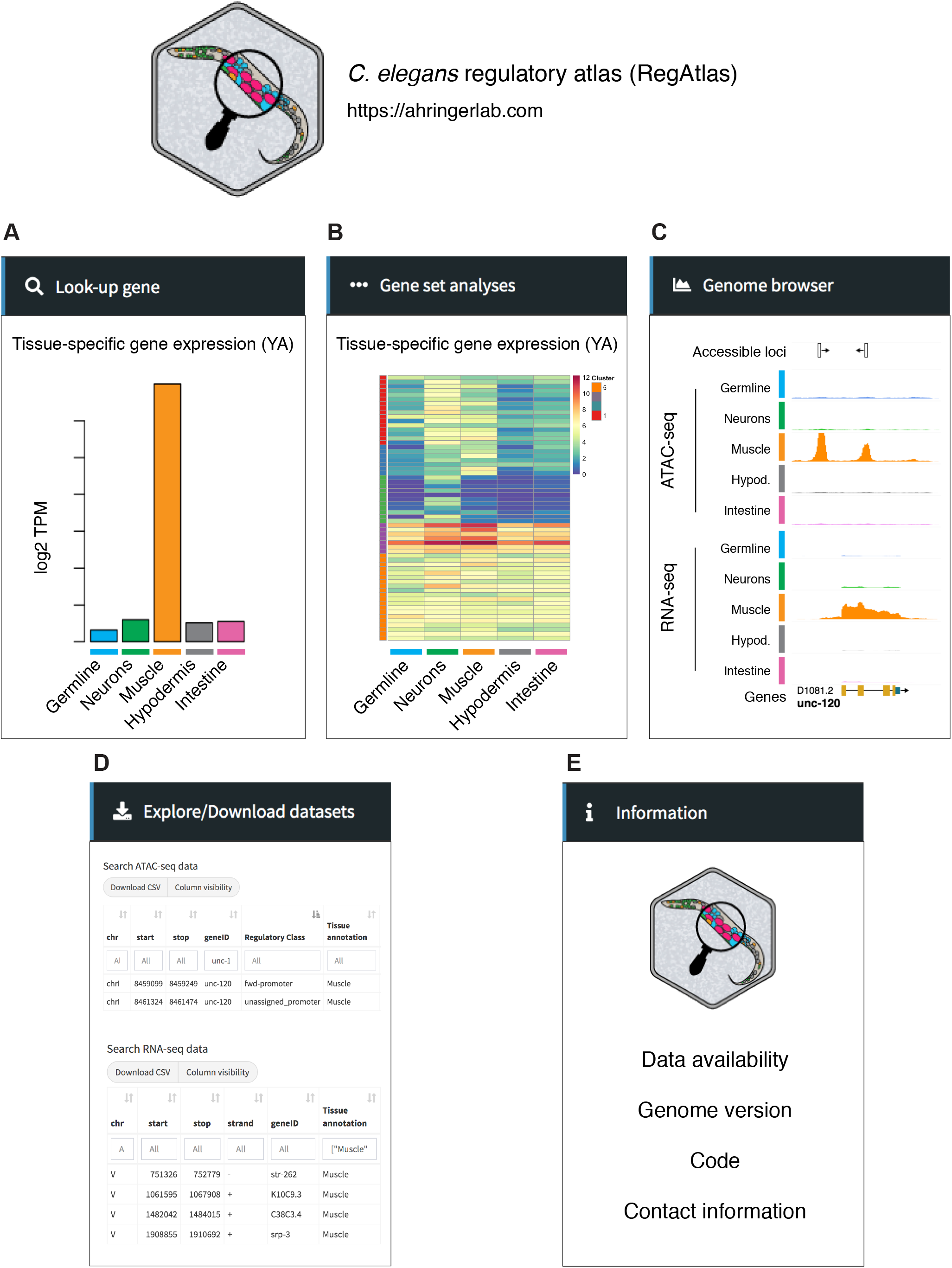
RegAtlas, a web interface to explore gene expression and chromatin accessibility datasets. Interface of RegAtlas, a web application developed to explore developmental and tissue-specific genomic datasets. RegAtlas is hosted at https://ahringerlab.com. Its use is entirely anonymous and performed queries are not saved. (A) Tab to query information on a single gene. (B) Tab to intersect a user-provided list of genes with tissue-specific and ubiquitous sets of genes defined in this study, visualize their expression across development or in adult tissues and perform GO enrichment analysis. (C) Tab to dynamically browse different types of genomic tracks (*e.g.* developmental or tissue-specific ATAC-seq and RNA-seq tracks) using an integrated JBrowse genome browser (Buels et al., 2016). (D) Tab to explore and download all processed datasets in tables. (E) An information tab is also available to get more details about the web portal.

